# Wild-captive contrasts in non-vocal communicative repertoires and functional specificity in orang-utans

**DOI:** 10.1101/2021.01.19.426493

**Authors:** Marlen Fröhlich, Natasha Bartolotta, Caroline Fryns, Colin Wagner, Laurene Momon, Marvin Jaffrezic, Tatang Mitra Setia, Caroline Schuppli, Maria van Noordwijk, Carel P. van Schaik

## Abstract

The creation of novel communicative acts is an essential element of human language. Although some research suggests the presence of this ability in great apes, this claim remains controversial. Here, we use orang-utans (*Pongo* spp.) to systematically assess the effect of the wild-captive contrast on the repertoire size of communicative acts. We find that individual communicative repertoires are significantly larger in captive compared to wild settings, irrespective of species, age-sex class or sampling effort. Twenty percent of the orang-utan repertoire in captivity were not observed in the wild. In Sumatran orang-utans, the more sociable species, functional specificity was also higher in captive versus wild settings. We thus conclude that orang-utans, when exposed to a more sociable and terrestrial lifestyle, have the behavioural plasticity to invent new communicative behaviours that are highly functionally specific. This productive capacity by great apes is a major prerequisite for the evolution of language and seems to be ancestral in the hominid lineage.

## Introduction

One of the core features of human language is its productivity, referring to the idea that signallers can create and understand novel utterances with novel meanings ^1^. While several other building blocks of language, including intentionality, reference and compositional syntax, have increasingly been documented among a wide range of non-human species ^2, 3, 4^, productivity seems to be very rare in the animal kingdom. Instead, animal communication systems are thought to rely heavily on evolved signals, that is, communicative acts whose form and function evolved under the influence of the effect on the recipient ^5, 6^. The current consensus is that all facial, vocal and most gestural expressions of non-human species have evolved through natural selection over long periods of time, and have become innate: the ability to produce them arises spontaneously during ontogeny, whereas its use is often fine-tuned by practice ^7, 8, 9^.

Recent research on great apes, our closest living relatives, suggests that it may be timely to distinguish between innate animal signals and those that are acquired developmentally ^10^. Great apes have provided most comparative evidence for the cognitive building blocks and selective pressures shaping the human communication system ^11, 12, 13^. But recent work has also shown that some of their gestures and sounds are apparently innovated and maintained over time ^14, 15, 16, 17, 18, 19, 20^, and play the same role in the communication process as evolved signals do – we could thus call them invented signals. Because targeted studies to estimate the extent of productivity in great apes are so far lacking, we here examine this question by comparing the same species in the wild and in a novel setting, captivity ^10, 21^. The contrast allows a direct test of how repertoires respond to the changes in the socio-ecological environment. In captivity, individuals face less competition for food, have more spare time, are closer together, are more often visible to each other, and more on the ground than in the wild ^22, 23^. Especially in fission-fusion species such as orang-utans (*Pongo* spp.), interaction rates in contexts such as social play, grooming, conflict situations and mating are boosted in captive settings e.g. ^24, 25, 26^, which may favour the production of innovative communicative acts, and thus cause differences in the communicative repertoires of individuals and groups.

The captive-wild contrast also allows us to examine the extent to which the meaning of communicative acts depends on the context in which they are used. We predict that the learned, “species-atypical” communicative acts have high functional specificity, i.e. a highly context-specific production ^10, 27, 28^, a phenomenon reflecting communicative plasticity. This is because the use of invented signals inevitably implies intentionality: naïve individuals observed and learned them,inferred their meaning from context and reactions, and subsequently used them in the same context with the same function as their original inventor but see ^29, 30^ for contrary views on highly conserved signal production. In contrast, evolved signals may be used intentionally or non-intentionally. They may therefore vary in functional specificity, and show more communicative flexibility: the same communicative act serves several different goals or functions, relying on context to provide disambiguation ^2, 31^.

The aim of the present study was to examine repertoires and functional specificity of close-range communicative acts, in both wild and captive populations of orang-utans, the great ape genus which is in our view ideal for this avenue of research. Systematic studies on the gestural repertoire of captive orang-utans have demonstrated that their propensity for elaborate and flexible gesture use parallels that of other great apes ^32, 33, 34^. This suggests that social propensities can be fully expressed in captivity, as individuals do not need to be solitary in order to obtain sufficient food, whereas in the wild they may have fewer interaction opportunities and communication is hampered by arboreality and obscuring vegetation. In addition, mothers are the predominant communication partner of infant orang-utans in the wild ^35, 36^, so that there is a limited need for the production of extensive communicative repertoires. There are no systematic wild-captive comparisons of apes’ communicative behaviour to date, but we assume that contrasts must be larger for orang-utans than any other great ape taxon.

We examined non-vocal (i.e. gestural and facial) communicative acts of Bornean and Sumatran orang-utans (*Pongo pygmaeus/abelii*) in two wild populations and five zoos. In a first step, we established the repertoires and functions (presumed goals of communicative acts, with outcomes that apparently satisfied the signaller) of orang-utans’ non-vocal communicative acts, building on previous work conducted on chimpanzees and captive orang-utans ^33, 37^, but separately for wild and captive settings. We then tested several predictions about how setting affected individual repertoire sizes and functional specificity of signal types, while controlling for important confounding variables such as age-sex class and sampling effort. First, captivity should result in larger communicative repertoires because of boosted territoriality, sociability and interaction rates. As a result, wild repertoires should be a subset of the captive ones, except for those communicative acts that cannot be expressed in captivity (“wild-only”). Second, we expect that the form of these communicative acts expressed only in captivity should be tightly linked to the more terrestrial lifestyle or the increased sociability, especially in the less terrestrial Sumatran orang-utans. Third, the “invention” of additional signals in captivity should be accompanied by a wild-captive contrast in functional specificity. We expect to find this contrast especially or even exclusively in Sumatran orang-utans, because sociability and interaction rates are reportedly higher in the Northwest-Sumatran population compared to the Bornean populations ^38, 39^. The effect of this setting-species interaction on functional specificity will allow us to derive important conclusions on the plasticity and flexibility underlying communicative interaction in the *Pongo* genus.

## Results

### Communicative acts across settings

A total of 40 distinct signal types were identified across all settings, out of which 34 were observed in Bornean (captive: N = 27, wild: N = 24) and 39 in Sumatran orang-utans (captive: N = 37, wild: N = 32). Plotting the cumulative number of identified communicative acts over the course of the observation period indicated that study groups have been sufficiently sampled to grasp complete repertoires (Fig. 1, S1, S2, S3), except for two captive groups of Bornean orang-utans (Apenheul and Cologne, see Fig. S1). In Table S1 we provide definitions for all coded behaviours and their relation to previous work on orang-utans’ communicative repertoire. The majority of signal types (N) and cases (n) consisted of manual (N = 19, n = 5106) and bodily signals (N = 18, n = 2212), whereas considerably fewer facial acts (N = 3, n = 110) act were observed (see Tab. S2 for detailed overview of signal presence in relation to settings, species, subjects and age classes). The relatively small repertoire of facial signals may be partly due to our strict criteria of inclusion into the repertoire (see methods).

**Fig. 1.**
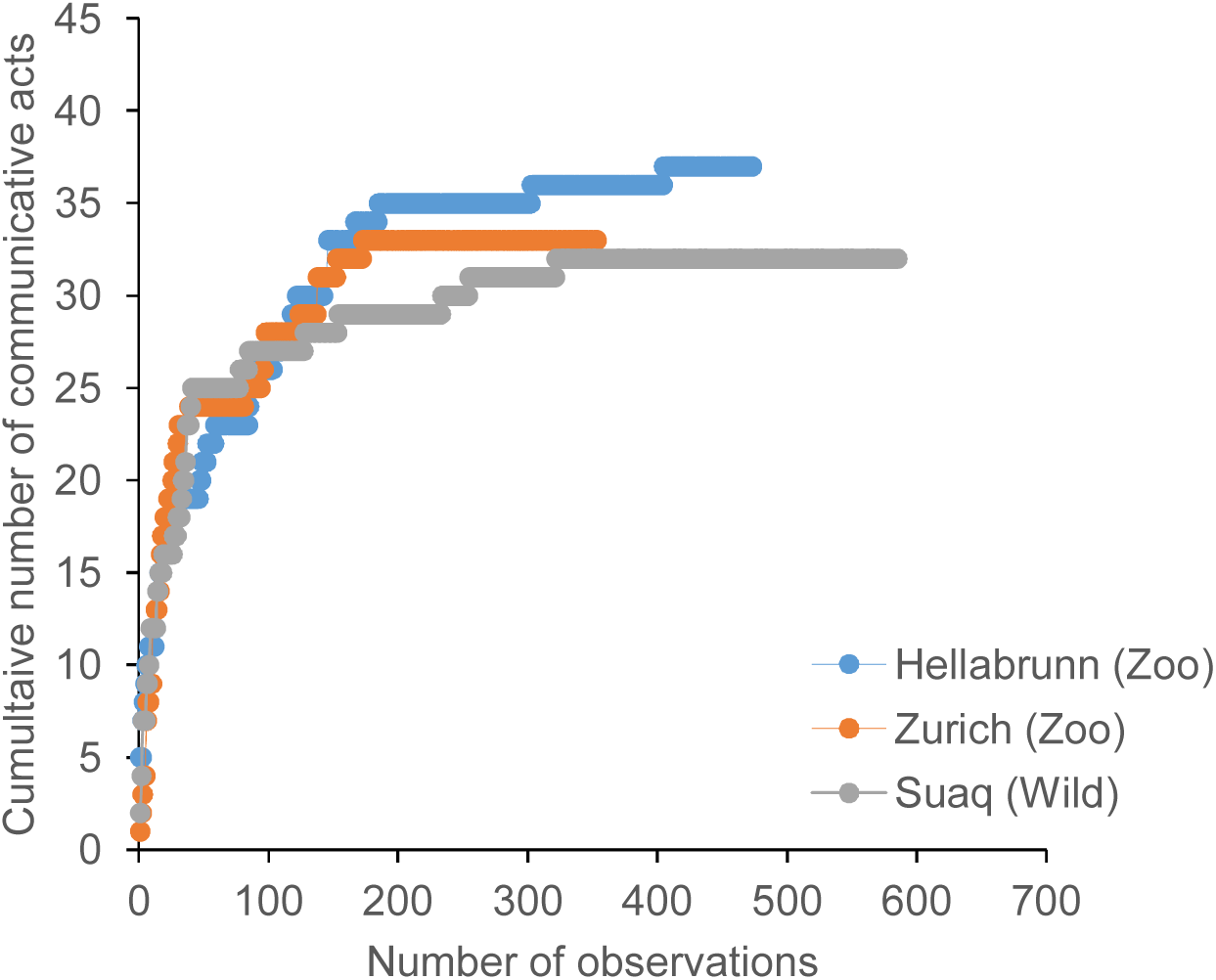
Cumulative number of identified communicative acts over observation time. Asymptotes are depicted separately for the captive subjects (Hellabrunn, Zurich) and wild populations (Suaq) of Sumatran orang-utans in this study.

We first tested the first prediction that captivity should result in enlarged communicative repertoires at the aggregate level owing to boosted sociability and interaction rates. We found that the majority of communicative acts (N = 27) was shared across orangutan species and research settings, but that 9 communicative acts were restricted to captivity (e.g. “roll on back”, “throw object”, “somersault”, see Fig. 2), and 2 to the wild (“loud scratch”, “shake object”, see Fig. 2), thus confirming the prediction. Out of these, seven captivity-specific and one wild-specific acts were observed in the Sumatran species only (e.g. “rub on body”, “head-butt”, “shake object”), whereas one behaviour (“spin”) was observed in captive Borneans only (see Tab. S2 for a detailed overview communicative acts across settings and species). A more conservative way of testing the prediction is by producing a list based on all previous studies. We found that three of the communicative acts we found only in captive settings (“throw object”, “rise up”) or wild settings (“shake object”), were also observed in other species-setting combinations in other studies, which leaves at least seven captivity-only acts and one wild-only act (see Tab. S3, note that Cartmill & Byrne [2010] do not specify which gestures were observed in which orangutan species). This more conservative test thus also confirms the prediction.

**Fig. 2.**
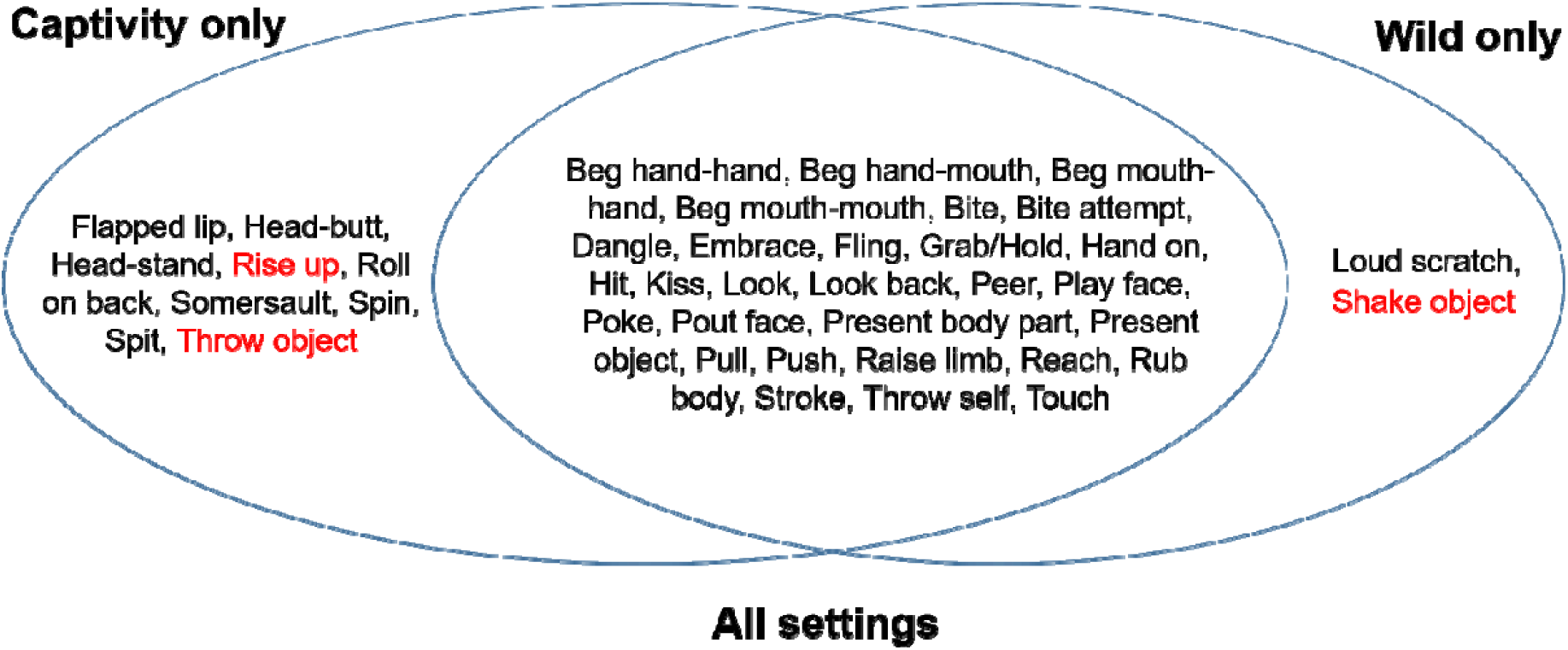
Overview of communicative acts observed only in different research settings. Communicative acts observed in the contrasting setting by other studies are marked in red.

To ensure that these differences between captivity and the wild do not reflect differences in social opportunities (e.g. with regard to the availability same-age play partners), we compiled separate play repertoires for mother-offspring (for which there is no change in partner availability between natural and captive settings) versus same-aged interactions (Tab. S4). A graphical analysis revealed no substantial difference between captive versus wild play repertoires with regard to mother-offspring interactions (25 vs 24 in Sumatrans, 22 vs 18 in Borneans). For peer play interactions, repertoire sizes apparently differed between settings for Sumatrans (26 vs. 19), but not Borneans (12 vs. 11). Differences in repertoire sizes between captive and wild settings are thus not driven by partner availability alone. Given that 13 communicative acts used in same-aged play interactions were exclusively used in Sumatrans (Tab. S4), we suggest that repertoire size is driven more by social opportunities in Sumatrans than in Borneans.

### Individual repertoire sizes across settings

The average Bornean orang-utans had 1.9 more signal types in its repertoire (captive: mean ± SD = 8.1 ± 3.8, wild: 10 ± 4.7), while the average Sumatran had 6.8 more signal types (captive: mean ± SD = 15.3 ± 6.5, wild: 8.5 ± 7.2), with a larger between-individual variation. With regard to age classes, we found that younger immatures had the largest repertoire on average (mean ± SD = 14.8 ± 5.9, N = 42), followed by older immatures (10.1 ± 8, N = 41), and adults (mean ± SD = 7.3 ± 3.9, N = 37).

Using a linear mixed model (LMM), we tested how setting, species and confounding variables such as sex, age class and sampling effort affected the number of communicative acts in individuals recorded during the study (for details see methods). The full model including the key test predictors (i.e. setting and species) fitted the data better than the null models irrespective of the subsets used (LRT all individuals: χ^2^_3_ = 14.058, *P* = 0.003, *N* = 70; individuals with > 50 interactions: χ^2^_3_ = 15.131, *P* = 0.002, *N* = 44). As expected, the number of communicative acts were strongly affected by the number of samples contributed to the dataset (see Tab. 1 for output of the model using the restricted dataset, Tab. S5 for the model including all individuals). Irrespective of the effects of sampling effort, however, we found that captive individuals deployed a significantly larger variety of communicative acts than their wild counterparts, again confirming our prediction. We also found that individual repertoires of Sumatran individuals exceeded those of their Bornean counterparts (after removing the non-significant interaction term, Tab. 1 a, Fig. 3 a, b). In addition, younger individuals produced significantly more different communicative acts than adults (Tab. 1 a, Fig. S4). These results are consistent with our expectations that young individuals (i.e. all immatures, but especially those below the age of 5 years) regularly use a larger communicative set than adults, particularly for the function of soliciting social play and food sharing. This is further supported by descriptive results on presumed goals and outcomes of communicative acts (see ESM, Tab. S6, S7) suggesting that interactions in both affiliative and conflict situations rather than co-locomotion or food-sharing underlie the proliferation of communicative acts in captive settings.

**Tab 1.**
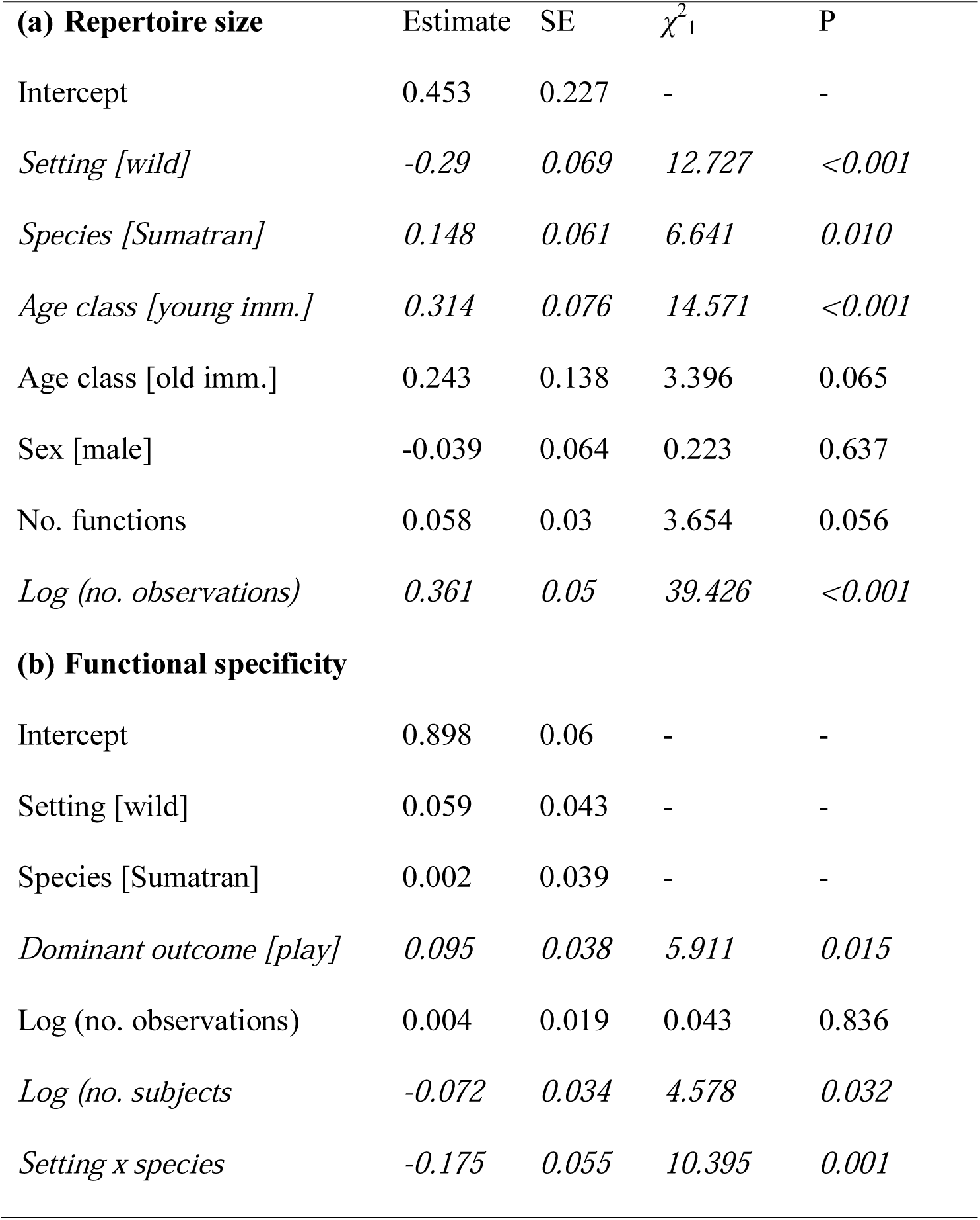
Effects of research setting, orang-utan species, and control variables on (a) repertoire size of individuals (N = 44), and (b) functional specificity of communicative acts (N = 114), derived using LMMs with a Gaussian error structure and identity link function. Significant effects (*P* < 0.05) are depicted in italics.

**Fig. 3.**
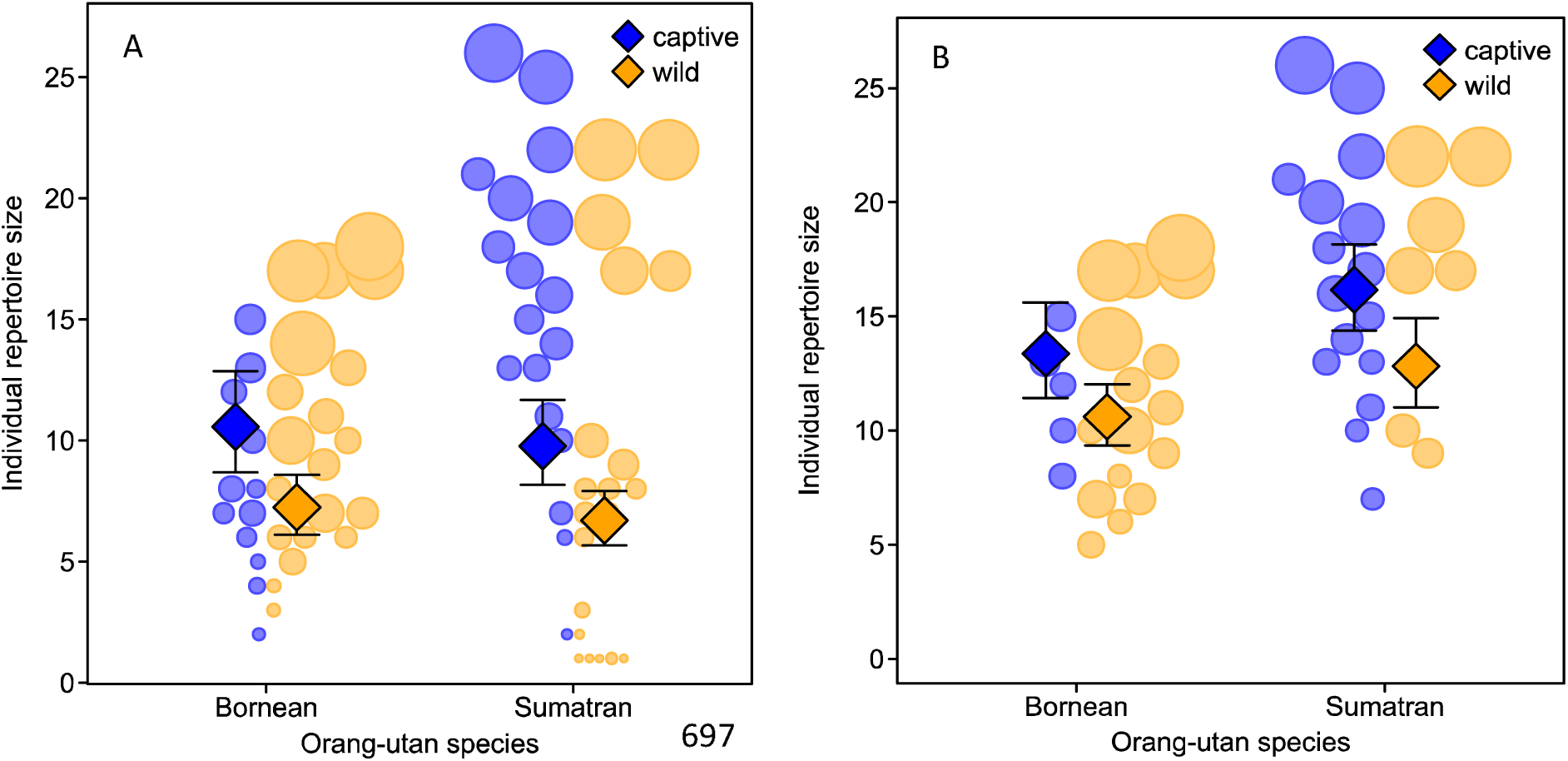
Number of observed types of communicative acts per individual as a function of research setting and orang-utan species, with **(A)** all subjects included, and **(B)** restricted to subjects with > 50 samples. Circles represent different individuals with area corresponding to sample size, Diamonds depict model estimates with 95% confidence intervals (all other variables centered to a mean of zero).

### Functional specificity of communicative acts across settings

We systematically tested the second prediction on functional specificity of communicative acts depends on research setting and orang-utan species using a LMM, which also included confounding variables such as outcome and sampling effort. The full model including the key test predictors (i.e. setting, species, dominant outcome) fitted the data better than the null models (LRT: χ^2^ _3_ = 22.612, *P* < 0.001, *N* = 114). There was a significant interaction between research setting and orang-utan species: while specificity scores in captivity did not differ between the species, we found a significantly lower functional specificity in wild Sumatrans compared to their captive counterparts (Fig. 4). Irrespective of this result, specificity was significantly higher for communicative acts predominantly used to solicit play and observed in a larger number of subjects (Tab. 1 b). Our findings thus support the prediction that larger repertoire sizes in captivity should be accompanied by an increase in average functional specificity.

**Fig. 4.**
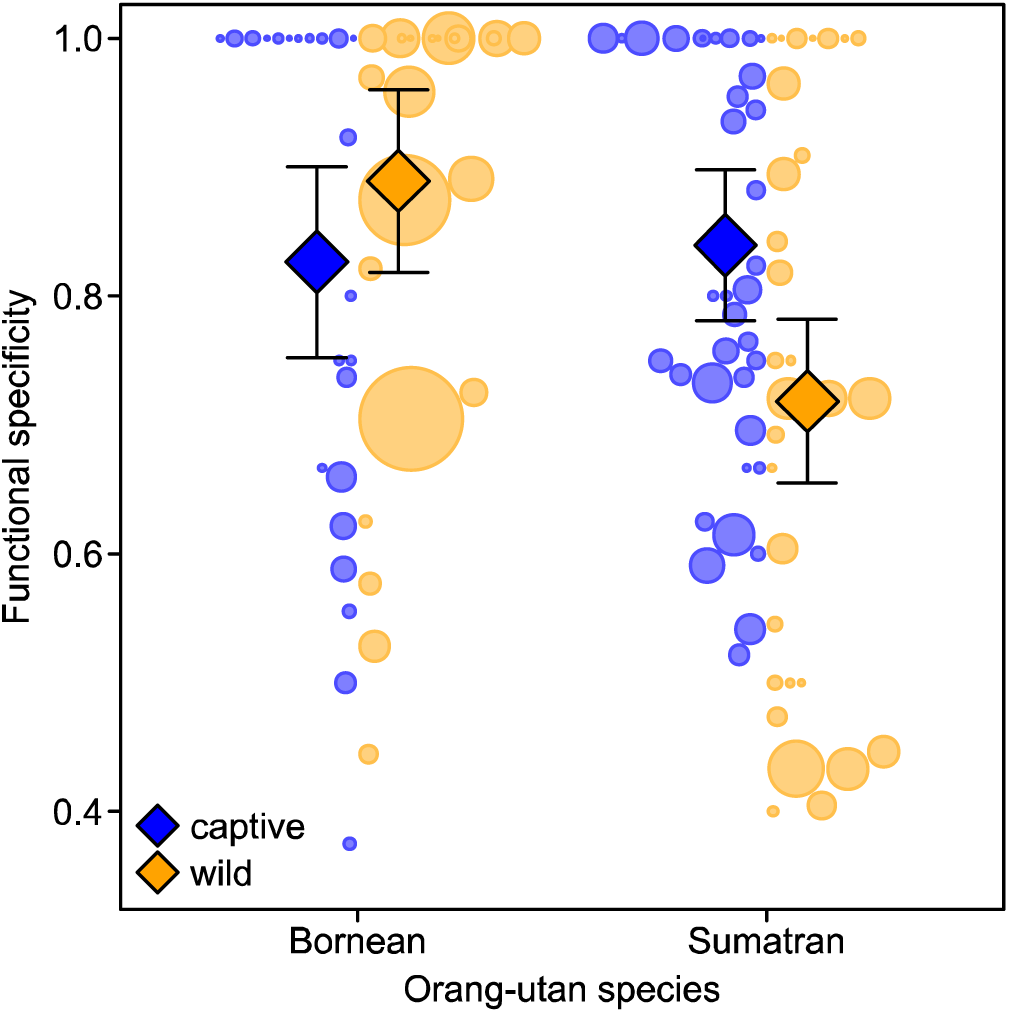
Functional specificity of communicative acts as a function of research setting and orangutan species. Circles represent different signal types with area corresponding to sample size, diamonds depict model estimates with 95% confidence intervals (all other variables centered to a mean of zero).

## Discussion

Answering the question whether our primate relatives possess the behavioural plasticity, or creative capacity, to complement their species-typical repertoires by inventing novel signals from scratch is highly relevant to theories of language evolution. Yet, to date no study explicitly and systematically examined communication systems of apes exposed to novel socioecological conditions relative to the wild baseline situation. Here, we adopted a 2 × 2 comparative design, investigating repertoire sizes and functional specificity of communicative acts in zoo-housed and wild groups of two different orang-utan species with different sociability and terrestriality. By examining the captive-wild contrast in these two related species we tested the prediction that captive environments favour the emergence of novel communicative acts, which should also have higher functional specificity. Moreover, comparing species differences related to differential sociability and terrestriality on one hand, and setting on the other, we expected that Sumatran orang-utans, but not Borneans, would show a wild-captive contrast in average functional specificity, offering insights into the degrees of plasticity and flexibility underlying the communicative repertoires of orang-utans.

Consistent with our first prediction, communicative repertoires on both the aggregate and individual level were larger in captivity as compared to the wild, even after controlling for the expected effects of age class and sampling effort (i.e. irrespective of whether all or only highly sampled individuals are included in the analysis). There may be some doubt that a single study can exhaustively sample signal repertoires. We therefore also compared the captive-wild contrast for each species using all available studies. This comparison supported the conclusions based on our study, in that the actual repertoire *composition* found in previous studies in the same setting and species revealed no major differences (Tab. S2, S3). First, our own findings regarding zoo repertoires (i.e. 27 different communicative acts in captive Borneans and 37 in captive Sumatrans, as compared to 24 and 32 communicative acts in the wild, respectively) are broadly consistent with the available systematic studies in single settings. Liebal and colleagues ^32^, studying two captive groups of Sumatran orang-utans, reported a repertoire of 34 signal types (29 gestures, and 5 facial expressions). Like them, we found that the majority of communicative acts were used to solicit social play and food transfers. Cartmill & Byrne ^33^, examining two zoo groups of Bornean and one group of Sumatran orang-utans, identified 38 types of gesture and facial expressions that allowed the analysis of “intentional meaning”. Second, the first systematic study on mother-offspring gesture use among wild orang-utans, conducted at the Bornean population of Sabangau Forest, identified 21 gesture types that met the criteria for inclusion into the repertoire ^35^. With 24 different observed communicative acts observed in our study population at Tuanan, it seems like the inclusion of communicative interactions outside the mother-offspring bond does not result in a substantially larger repertoire size. Thus, we can conclude that moving wild orang-utans into captivity leads to a 20 to 25% increase (i.e. 7 to 9 acts “gained”, 1 to 2 acts “lost”) in their repertoire of communicative acts.

The second prediction we made was that the form of these communicative acts expressed only in captivity should be tightly linked to the increased sociability and more terrestrial lifestyle. As expected, differences in repertoire size were particularly pronounced for presumed goals related to seeking body contact (“Play/affiliate”, “Groom”, “Sexual contact”) and social conflict (“Move away” and “Stop action”). Interaction rates with these outcomes are greatly boosted in captivity, where a more differentiated use of bodily communication is both enabled and required ^10^. Captive facilities are stable, plentiful and predator-free environments that may provide opportunities for, and even require (e.g. due to increasing conflict with limited space) signal inventions and innovations, just like they foster innovations in general ^40, 41^. Our findings thus provide direct evidence that the new environments we have created for great apes boost the invention of new signals, which may spread through social learning. Indeed, captive settings in general have generated extensive and convincing evidence for invented (“species-atypical”) signalling, encompassing novel pant-hoot variants ^42^, and “whistling” ^15^, as well as pointing with hands and fingers ^43, 44^, “raspberries” and “extended grunts” in chimpanzees ^16^. Although detailed captive-wild comparisons are, to our knowledge, so far lacking for other great apes, wild-captive contrasts are probably larger in orang-utans than any other great ape taxon. In conclusion, our results support the notion that the new opportunities and needs linked to captivity may lead to a proliferation of signal invention.

We also expected that the form of these communicative acts expressed only in captivity are linked to a more terrestrial lifestyle. Indeed, we found that the communicative acts that are exclusively (or overwhelmingly) produced in captive settings are strongly linked to the more terrestrial nature of their artificial habitat: “somersaults”, “spitting”, “head-stands”, communicative acts that involve either the ground or objects obtained from the ground, would be very difficult to perform by wild orang-utans with their purely (Sumatra) or predominantly Borneo: ^45^ arboreal lifestyle. This setting effect is not attributable to the presence of certain interaction partners alone: by comparing the repertoires for mother-offspring and peer play interactions we demonstrated that differences between interaction dyads with regard to wild-captive contrasts were only strong in Sumatran orang-utans. It thus appears that the new affordances of captive settings, on top of the elevated exposure to certain social contexts, enabled orang-utans to better exploit their (communicative) motion spectrum, resulting in novel communicative movements that may independently and predictably such as spitting as an attention-getter, see ^46^ be invented in several captive colonies and species. This also confirms earlier reports making the case that the complex individual-based fission fusion structure of orang-utans and their sophisticated social-cognitive skills seem to be reflected in a highly variable communicative repertoire ^32, 47^, illustrating their remarkable behavioural plasticity.

Finally, we predicted that the additional signals in captivity should be accompanied by an increase in functional specificity. In line with our predictions, we found that Sumatrans in the wild exhibited a lower average functional specificity compared to their captive counterparts, while average functional specificity in captivity did not differ between the species. When comparing Sumatrans’ communicative acts used only in captivity (“invented signals”) with those used in both research settings, captivity-only acts appeared to be on average more functionally specific (although the small sample prevented inferential analyses). In other words, wild Sumatrans seem to use their communicative acts more flexibly (i.e. redundantly) across presumed goals than in captivity, which appears to be largely due to captive Sumatrans’ use of invented context-dependent, and therefore functionally specific acts (i.e. those that have “tight meanings” according to ^33^) not present in the wild repertoire. In contrast, the relatively low interactions rates (due to few social opportunities) of wild Bornean orang-utans ^38, 39, 48^ seem to be reflected in a lesser need to use signals flexibly across contexts. Together, these results corroborate our expectation that average functional specificity increased with repertoire size in captive Sumatran, though not Bornean orang-utans.

To convey a message that can be understood by a targeted recipient, an intentional agent may use two alternative “strategies”. First, she can use one and the same communicative act for several different functions, relying on context or other information (e.g. possibly age difference or sex relative to recipient) to disambiguate between ambiguous meanings. This “flexible” communicative strategy produces redundancy in the communicative repertoire. Alternatively, she can use a communicative behaviour that is specific to one interaction outcome – and, if not available, invent one for which naïve recipients (over repeated instances of interaction) infer their meaning from context and reactions, which they may subsequently use in exactly the same contexts with exactly the same function as their original inventor. This “plastic” communicative strategy produces productivity in the communicative repertoire. Sumatran orang-utans in the wild seem to rely somewhat more on the flexibility option; their captive counterparts, however, rely more on the plasticity option because repeated interactions with the same partners in captivity allow them to establish novel signal meanings more rapidly. This outcome is expected because novel signals are most likely to be understood and thus maintained when they are highly context-specific.

We argue that distinguishing species-typical from invented communicative acts matters greatly for current debates on language origins see also ^10^. Language reflects extreme communicative plasticity. Words that make up language are predominantly invented noises used intentionally, and the productivity feature of language fundamentally relies on the ability to produce such new noises, which are used intentionally and rapidly acquire a shared meaning due to their high context-specificity. Functionally specific signals are often more effective and efficient in complex interactional exchanges (including language), because they are less dependent on context and thus less ambiguous. Reliance on context may be less favourable with increasing message complexity (multiple intertwining messages), because it increases both (i) the risk of misunderstanding, and (ii) the “decoding work” necessary for the recipient, and thus the risk that the signal is ignored.

Although our knowledge of the taxonomic distribution of invented signals is still incomplete, they are so far reported almost exclusively for great apes. Ongoing work supports this pattern, in that great apes are increasingly documented to make up new vocal e.g. ^15, 16, 49^ and gestural e.g. ^18, 50^ ‘signals’ in the novel conditions of captivity, whereas reports from other taxa are rare but see e.g. ^51, 52, 53, 54^. Such communicative creativity may therefore be most common in, or even limited to, species with intentional communication. These findings imply that once the conditions were in place that favoured the open-ended use of invented expressions, our hominin ancestors readily responded to this opportunity, because they could build on a long evolutionary history of communicative creativity. This might explain why language evolved in the hominin lineage and not others that found themselves in similar conditions (e.g. those canids that also rely on interdependent foraging and cooperative breeding).

## Materials and Methods

### Data collection

Data were collected at two field sites and five captive facilities (zoos). We observed wild orang-utans at the long-term research sites of *Suaq Balimbing* (03°02’N; 97°25’E, Gunung Leuser National Park, South Aceh, Indonesia) and *Tuanan* (02°15’S; 114°44’E, Mawas Reserve, Central Kalimantan, Indonesia), on a population of wild Sumatran (*Pongo abelii*) and Bornean orang-utans (*Pongo pygmaeus wurmbii*), respectively. Both study sites consist mainly of peat swamp forest and show high orang-utan densities, with 7 individuals per km^2^ at *Suaq* and 4 at *Tuanan* ^55, 56^. Captive Bornean orang-utans were observed at the zoos of Cologne and Münster, and at Apenheul (Apeldoorn), while Sumatran orang-utans were observed at the zoo of Zurich and at Hellabrunn (Munich; see EEP studbook for details on captive groups; Becker 2016). While captive Sumatran orang-utans were housed in groups of nine individuals each, captive Bornean groups were smaller (on average four individuals, with the one in Apenheul including only a mother and her dependent and independent offspring). Signallers included in this study consisted of 33 Bornean (21 wild/12 captive) and 38 Sumatran orang-utans (20 wild/18 captive; see Tab. S8 for detailed information on subjects and group compositions).

Focal observations were conducted between November 2017 and October 2018 (Suaq Balimbing: November 2017 – October 2018; Tuanan: January 2018 – July 2018, European zoos: January 2018 – June 2018). At the two field sites, these observations consisted of full (nest-to-nest) or partial follows (e.g. nest-to-lost or found-to-nest) of mother-infant units, whereas in zoos 6-hour focal follows were conducted. Two different behavioural sampling methods were combined: First, presumable intra-specific communicative interactions of all observed social interactions of the focal either as signaller or receiver with all partners (*N* = 7137 acts), and among other conspecifics (*N* = 888 acts) present were recorded using a digital High-Definition camera (Panasonic HC-VXF 999 or Canon Legria HF M41) with an external directional microphone (Sennheiser MKE600 or ME66/K6), enabling recordings of high-quality footage. In captive settings with glass barriers, we also used a Zoom H1 Handy recorder that was placed in background areas of the enclosure whenever possible. Second, using instantaneous scan sampling at ten-minute intervals, we recorded complementary data on the activity of the focal individual, the distance and identity of all association partners, and in case of social interactions the interaction partner as well as several other parameters. During ca. 1600 hours of focal observations, we video-recorded more than 6300 communicative interactions.

### Coding procedure

A total of 2655 high-quality recordings of orang-utan interactions (wild: 1643, captive: 1012) were coded using the program BORIS version 7.0.4. ^58^. We designed a coding scheme to enable the analysis of presumably communicative acts directed at conspecifics (i.e. behaviours that apparently served to elicit a behavioural change in the recipient and were mechanically ineffective i.e. achieve a presumed goal without physical force; see also ^33^. Manual, bodily, and facial acts were defined and aligned (see Tab. S1) based on previous studies on orang-utan communication in captive ^26, 32, 33, 46^ and wild settings ^35, 36, 59, 60^. Comparing our dataset to this literature, we then identified the subset of setting- and species-specific communicative acts. Although we also coded vocalizations based on field studies ^61^, we did not include vocalizations in the analyses of repertoire and functional specificity as we could not equally pick up soft, low-frequency sounds in captive and wild settings, which hampered the fine-grained comparison across settings. For each communicative act, we coded the following “modifiers”: presumed goal following the distinction of ^33^, outcome, and other variables not directly relevant in this study (see Tab. S9 for levels and definitions of all coded variables). To ensure inter-observer reliability, we evaluated the coding performance of all observers with alternating datasets using the Cohen’s Kappa coefficient ^62^ during an initial training period and at regular intervals afterwards. Trained observers (MF, NB, CF, CW, LM, MJ) proceeded with video coding only if at least a ‘good’ level (κ _>_ 0.75) of agreement was found for signal type, articulator, sensory modality, context, and response. For our repertoire analyses, we plotted the cumulative number of communicative behaviours over the number of coded interactions for each study group (Fig. S1, S2) and for a subset of highly sampled individuals (Fig. S3), to estimate how many observations are necessary to grasp the repertoire of these groups/individuals as indicated by an asymptote. Communicative acts were counted as part of individuals’ repertoire only when observed at least twice per subject.

To analyse functional specificity, we focused on goal-outcome matches ^33^ or apparently satisfactory outcomes ^37^, that is, whether the interaction outcome aligned with presumed goals identified by observers. We considered only those signal types that were produced at least three times towards a particular interaction outcome cf. ^37^. We defined functional specificity depending on how often a communicative act was produced towards an apparently satisfactory outcome (ASO), adopting the definitions of Cartmill and Byrne ^33^. Signal types that were used mainly towards a single interaction outcome, i.e. more than 70% of the time, were defined as having “tight meanings”. Signal types used frequently towards a single ASO, i.e. 50–70% of the time, were defined as having “loose meanings”. Finally, signal types that were used less than 50% of the time towards a single ASO were classified as having “ambiguous meanings”. For instance, “somersault” was exclusively produced to initiate “Play/affiliate” interactions (specificity value of 1, tight meaning), whereas “touch” was produced towards several different interaction outcomes, e.g. “Play/affiliate”, “Share food/object” and “Co-locomote” (specificity values < 0.7, ambiguous or loose meaning).

### Statistical analyses

We ran two separate linear mixed models LMMs; ^63^ with a Gaussian error structure and identity link function to examine sources of variation in (a) individual communicative repertoires (i.e. number of signal types used at least twice) and (b) specificity in signal function. We used LMMs rather than GLMMs in this study because it has recently been shown that linear models are more robust to violations of distributional assumptions ^64, 65^. We ran model (a) for two subsets of our data: first, including all individuals regardless of sample size; second, including only those individuals that contributed more than 50 communicative acts to the dataset (as the graphical inspection of asymptote plots in Fig. S3 suggested that this is a representative number to estimate individual repertoire sizes).

In model (a), which had individual repertoire size as response variable, we included research setting (2 levels: captive, wild) and orang-utan species (2 levels: Bornean, Sumatran) as our key test predictors. Because we assumed that the effect of research setting might depend on genetic predisposition (i.e. species), we included the interaction between these two variables into our model. Moreover, immature individuals often contribute the majority of both signal cases and types ^2^, for reviews see ^31^ and the composition of study groups differed with regard to age-sex classes, hence we made sure that age class was taken into account in our analyses. We included the following variables as additional fixed effects (control predictors) into the models: subjects’ age class (3 levels: “adult”: females > 15 years, males > 16 years; “older immature”: independent and dependent immature > 5 years of age, “younger immature”: dependent immature < 5 years of age), sex (2 levels: female, male), the number of interaction outcomes the subject communicated for at least twice (range = 1–6, only four outcomes were coded in captive Borneans), and the number of observations (range = 5– 467). To control for repeated measurements within the same sampling unit, group identity was treated as random effect. To keep type 1 error rates at the nominal level of 5%, we also included all relevant random slopes components within group ID ^66^.

In model (b), which had specificity in signal function as response variable, we included orang-utan species (two levels: Bornean, Sumatran), research setting (two levels: captive, wild) and dominant outcome (two levels: play, non-play), and the interaction between setting and species as our key test predictors. To control for confounding effects of sampling effort, the number of subjects contributing to the use of a signal type (range = 1–21) and the number of observations (range = 1–787) in the respective setting as additional fixed effects (i.e. control predictors). To control for repeated observations of the same signal types across settings, signal type was treated as random effect ^67^.

All models were implemented in R (v3.4.1, ^68^) using the function *glmer* of the package lme4 ^69^. To control for collinearity, we determined the Variance Inflation Factors VIF; ^70, 71^ from a model including only the fixed main effects using the function *vif* of the R package car ^72^. This revealed no collinearity issues (maximum VIF = 2.8). Prior to running the models, we log-transformed the response variables and the control variables relating to sampling effort (number of observations/subjects), to achieve an approximately symmetrical distribution and avoid influential cases.). To test the overall significance of our key test predictors ^73, 74^, we compared the full models with the respective null models comprising only the control predictors as well as all random effects using a likelihood ratio test ^75^. Tests of the individual fixed effects were derived using likelihood ratio tests (R function *drop1* with argument “test” set to “Chisq”).

## General

We thank Erin Vogel and Suci Utami Atmoko (Tuanan), Kerstin Bartesch (Tierpark Hellabrunn), Claudia Rudolf von Rohr (Zoo Zürich), Alexander Sliwa (Kölner Zoo), Simone Sheka (Allwetterzoo Münster) and Thomas Bionda (Apenheul Primate Park) as well as all research staff and zoo keepers for a fruitful collaboration during this study. We gratefully acknowledge Clemens Becker for providing the EEP studbook, the Indonesian State Ministry for Research and Technology (RISTEK), the Indonesian Institute of Science (LIPI), the Directorate General of Natural Resources and Ecosystem Conservation – Ministry of Environment & Forestry of Indonesia (KSDAE-KLHK), the Ministry of Internal affairs, the Nature Conservation Agency of Central Kalimantan (BKSDA), the local governments in Central Kalimantan, the Kapuas Protection Forest Management Unit (KPHL), the Bornean Orang-utan Survival Foundation (BOSF) and MAWAS in Palangkaraya. Moreover, we thank Simone Pika and Eva Luef for providing some of the essential equipment, Santhosh Totagera for coding assistance, Uli Knief and Alex Hausmann for help with a customized jitter-plot function.

## Author contributions

MF and CvS conceived of the study. MF designed the project, collected, coded and analysed data. NB, CF, CW, LM, MJ helped to collect, curate and code data. TMS, CS, MvN and CvS provided resources. MF wrote the manuscript with inputs from CF, LM, CS, MvN and CvS. All authors approved the submission of the manuscript.

## Data and materials availability

All data needed to evaluate the conclusions in the paper and the R code can be found on GitHub: https://github.com/MarlenF/repertoire-orang.

