## Supplementary tables and figures for "Wild-captive contrasts in non-vocal communicative repertoires and functional specificity in orang-utans"

**Table S1** Non-vocal communicative acts observed in this and other studies on orang-utans in captivity (*1-6*) and the wild (*7-10*), and that met the criteria for inclusion of the contextual analyses.

| **Behaviour** | **Definition (this study)** | **Other studies in captivity** | **Other studies in the wild** |
| --- | --- | --- | --- |
| **Manual acts** |  |  |  |
| Beg hand-hand | Attempts to obtain food out of rec's hand with hand |  |  |
| Beg hand-mouth | Attempts to obtain food out of rec's mouth with hand |  |  |
| Embrace | Bring one or both arms around other individual | Embrace (5,6) | Embrace (7,8) |
| Fling | Flail limbs towards other ind (usually arm) | *Schlagintention* [hit intention] (1), Hit intention (5), wave arm (6),  Beckon (2), fake (2), swat (2) | Hit away (8), Fling (9) |
| Grab/hold | Grab or hold onto other ind's body part longer than 2 s | Hold tight (5), Grab (6), grasp (6) | Grasp (8) |
| Hand on | Making long (> 2s) with other ind's body, fingers not closed | Put hand on head (6), Cover (6) | Hand on (9) |
| Hit | Hard and quick contact with other ind's body (e.g. With wrists or palms) | Slapping (3), *Schlagen* [hit] (4), Slap (5), Hit (6) | Hitting (8) |
| Loud scratch | Large, audible scratches on own body while "waiting" for recipient |  | Scratching (8), Loud scratch (10) |
| Poke | Soft and quick contact with other ind's body (e.g. With fingers or toes) | Nudge (1), poke (6), tap (6) | Poke (9) |
| Pull | Pull on other ind's body part or object that other ind is holding | *Ziehen* [pull], Pull (5,6) | Grasp (8), Grab-pull (9) |
| Push | Push other ind's body part or object that other ind is holding | Push/dragging (2), Push (5,6), | Push (9) |
| Raise limb | Raise arm or leg above own body (vertically) | Arms up (2), raise arm (2) |  |
| Reach | Extend arm or leg towards other ind or object other ind is holding | Hand extension *(3), Handausstrecken* [hand extension] (4), Extend arm (5), reach (6) | Hold out hand (8), Reach palm/wrist (9) |
| Shake object | Rapid back-and-forth movement of an object held in hands | Shake object (5, 6) | Branch shaking (5), Object shake (9) |
| Stroke | Moving fingers repeatedly and gently over another ind's body |  | Stroke (9) |
| Throw object | Throw object in the direction of another ind | *Object werfen* [throw object] (4), Throw object (5) | Throwing (8), Throw object (9) |
| Touch | Making brief (<2s) light contact with other ind's body | *Anfassen* [touch] (4), Gentle touch (5), touch (6) | Touch (8), Touch other (9) |
| **Bodily acts** |  |  |  |
| Beg mouth-hand | Attempts to obtain food out of rec's hand with mouth | Food beg orally (6) |  |
| Beg mouth-mouth | Attempts to obtain food out of rec's mouth with mouth | Put face on face (5), Food beg orally(6) | Mouth-mouth contact (4) |
| Bite | Lips and teeth closing are closing around body parts of another ind | Biting hands/feet (3), Bite in hand (5), Bite (6) | Biting (4), Bite (5) |
| Bite attempt | Mock bite other individual, mouth and lips touching other but teeth are not closing | *Beißintention* [bite intention] (1), Bite intention (5), air bite (6) | Lunge (7), Mock bite (8), muzzle-pushing (8) |
| Dangle | Hang above or in front of other ind, by moving in the substrate (e.g. rope net, liana) | *Strampeln* [struggle] (4), Swing (5, 6), Dangle (6) | Dangle (5) |
| Head-butt | Push head repeatedly onto other ind's body | Head-butting (3), Wrestle head-first (5) |  |
| Head-stand | Look through own legs at other ind ("self-handicapping" gesture) | Head-stand (5,6) |  |
| Kiss | Make contact with other ind with face and/or mouth | Lip touch (5), Kiss (6), mouth (6) | Kiss (7), mouth-mouth contact (8) |
| Look | Gaze intently at other ind, > 2 seconds, more than 15 cm |  | Watch (4), Fixed gaze (4) |
| Look back | Gaze intently at other ind, while turned in opposite direction | *Scheinflucht* [pretend escape] (1,4), Look back (6) |  |
| Peer | Look intently at what other ind is doing, less than 15 cm away | Approach face (5), peer (6) | Look at mouth (8) |
| Present body part | Move specific body part in v field of other ind | Offer body part (5), Present body part (5,6) | Presenting (8) |
| Present object | Move held object (e.g. In mouth, hand) in v field of other ind | *Object vorzeigen* [present object] (1, 4), Present object (5), show (6) | Object in mouth (9) |
| Rise up | Stand up bipedally |  | “extended posture” (7), Posturing (7, 8) |
| Roll on back | Lie down in front of other ind | Roll on back (6) |  |
| Rub body | Rub genitals on body of other ind | Touch with genital region (5) |  |
| Somersault | Move forward so that the body rolls end over end, making a complete revolution. | Somersault (6) |  |
| Spin | Rotate one’s body approximately 360° |  |  |
| Spit | Spit previously collected water in the direction of another ind | *Spucken* [spit] (1) |  |
| Throw self | Bring own body against or on top of another ind | Anspringen, Fallenlassen [jump at] (4), Jump at (5) |  |
| **Facial acts** |  |  |  |
| Flapped lip | Fold lower lip upwards |  |  |
| Play face | Play face with lower teeth exposed | Spielgesicht [play face] (4), Relaxed open mouth (5), Play face (6) | Play face (5), Relaxed open-mouth (6) |
| Pout face | Lips are shaped like an "o" | Funnel face (3), Pout face (5), Pout (6) | Pout face (5), pout moan face, Silent-pout face (6) |

**Table S2** Use of communicative acts in relation to setting, species and age class (Ad = adult, Im = older immature, Dp = younger immature, FS = Share food/object, GR = Groom, JT = Co-locomote, PL = Play/affiliate, SX = Sexual contact, ST = stop action; blue: wild-only, red: captivity-only, green: Sumatran-only/both settings).

| **Comm. act** |  |  | **Age class** | **Dominant outcome** | **Bornean** | | **Sumatran** | |  |
| --- | --- | --- | --- | --- | --- | --- | --- | --- | --- |
|  | **Type** | **No. subjects** |  |  | **captive** | **wild** | **captive** | **wild** | **Total** |
| beg hand-hand | manual | 36 | Ad, Im, Dp | FS | (1) | 152 | 55 | 98 | 306 |
| beg hand-mouth | manual | 33 | Ad, Im, Dp | FS | 4 | 82 | 64 | 73 | 223 |
| beg mouth-hand | bodily | 27 | Ad, Im, Dp | FS |  | 69 | 76 | 28 | 173 |
| beg mouth-mouth | bodily | 30 | Ad, Im, Dp | FS | 4 | 38 | 136 | 24 | 202 |
| Bite | bodily | 47 | Ad, Im, Dp | PL | 22 | 264 | 54 | 8 | 348 |
| bite attempt | bodily | 50 | Ad, Im, Dp | PL | 11 | 141 | 61 | 24 | 237 |
| dangle | bodily | 33 | Ad, Im, Dp | PL | 18 | 36 | 49 | 65 | 168 |
| embrace | manual | 23 | Ad, Im, Dp | PL | (1) | 8 | 23 | 18 | 50 |
| **flapped lip** | facial | 6 | Ad, Im, Dp | ST |  |  | 11 |  | 11 |
| fling | manual | 19 | Ad, Im, Dp | PL |  | 4 | 31 | 37 | 72 |
| grab/hold | manual | 65 | Ad, Im, Dp | PL | 104 | 787 | 178 | 348 | 1417 |
| hand on | manual | 47 | Ad, Im, Dp | PL, FS, GR | 64 | 177 | 29 | 148 | 418 |
| **head-butt** | bodily | 10 | Ad, Im, Dp | PL |  |  | 29 | 3 | 32 |
| **head-stand** | bodily | 3 | Ad, Im, Dp | PL |  |  | 4 | (1) | 5 |
| hit | manual | 20 | Ad, Im, Dp | PL | 4 |  | 83 | 4 | 91 |
| kiss | bodily | 37 | Ad, Im, Dp | PL | 5 | 57 | 62 | 2 | 126 |
| look at | bodily | 53 | Ad, Im, Dp | PL | 26 | 40 | 72 | 210 | 348 |
| **look back at** | bodily | 11 | Ad, Im, Dp | PL |  |  | 26 | (1) | 27 |
| **loud scratch** | manual | 11 | Ad, Im, Dp | JT |  | 5 |  | 29 | 34 |
| peer | bodily | 38 | Ad, Im, Dp | FS | 11 | 39 | 223 | 88 | 361 |
| play face | facial | 26 | Ad, Im, Dp | PL | 7 | 10 | 46 | 20 | 83 |
| poke | manual | 23 | Ad, Im, Dp | PL |  | 36 | 87 | 3 | 126 |
| pout face | facial | 7 | Ad, Im, Dp | PL |  | 10 | 6 |  | 16 |
| present body part | manual | 36 | Ad, Im, Dp | PL, GR | 7 | 87 | 28 | 26 | 148 |
| present object | manual | 16 | Ad, Im, Dp | PL | 8 | (1) | 14 | 4 | 27 |
| pull | manual | 61 | Ad, Im, Dp | PL, FS | 55 | 110 | 222 | 161 | 548 |
| push | manual | 50 | Ad, Im, Dp | ST, PL | 26 | 42 | 95 | 115 | 278 |
| raise limb | manual | 28 | Ad, Im, Dp | PL | 28 |  | 14 | 10 | 52 |
| reach | manual | 53 | Ad, Im, Dp | PL, FS | 23 | 18 | 88 | 107 | 236 |
| **rise up** | bodily | 7 | Ad, Im, Dp | PL |  |  | 3 |  | 3 |
| **roll on back** | bodily | 7 | Ad, Im, Dp | PL | 3 |  | 2 |  | 5 |
| **rub body** | bodily | 12 | Ad, Im, Dp | SX |  |  | 60 | 27 | 87 |
| **shake object** | manual | 1 | Ad, Im, Dp | PL |  |  |  | 5 | 5 |
| **somersault** | bodily | 6 | Ad, Im | PL | 6 |  | 4 |  | 10 |
| **spin** | bodily | 3 | Im | PL | 3 |  |  |  | 3 |
| **spit** | bodily | 1 | Im | PL |  |  | 5 |  | 5 |
| stroke | manual | 11 | Ad, Im | PL | (1) |  | 13 | 5 | 19 |
| **throw object** | manual | 7 | Im | PL | (1) |  | 10 |  | 11 |
| throw self | bodily | 19 | Ad, Im | PL | 12 |  | 58 | 2 | 72 |
| touch | manual | 66 | Ad, Im | PL | 77 | 598 | 156 | 214 | 1045 |
| Total |  | 70 |  |  | 532 | 2811 | 2177 | 1908 | 7428 |

**Table S3** Occurrence of communicative acts in captive (*1-6*) and wild (*7-10*) orang-utan repertoires (X = this study). Communicative acts depicted in red colour represent mismatches between our and other studies.

| **Communicative act** | **Bornean** | | **Sumatran** | |
| --- | --- | --- | --- | --- |
|  | **captive** | **wild** | **captive** | **wild** |
| ***Manual acts*** |  |  |  |  |
| Beg hand-hand | X | X | X | X |
| Beg hand-mouth | X | X | X | X |
| Embrace | X, 6 | X, 7 | X, 5, 6 | X, 8 |
| Fling | 6 | X, 9 | X, 5, 6 | X, 8 |
| Grab/hold | X, 6 | X | X, 1, 5, 6 | X, 8 |
| Hand on | X, 6 | X, 9 | X, 5, 6 | X, |
| Hit | X, 6 |  | X, 2, 5, 6 | X, 8 |
| Loud scratch |  | X |  | X, 8 |
| Poke | X, 6 | X, 9 | X, 5, 6 | X |
| Pull | X, 6 | X, 9 | X, 2, 5, 6 | X, 8 |
| Push | X, 6 | X, 9 | X, 2, 5, 6 | X |
| Raise limb | X, 6 |  | X, 6 | X |
| Reach | X, 6 | X, 9 | X, 5, 6 | X, 8 |
| Shake object | 6 | 7, 9 | 1, 6 | X |
| Stroke | X | 9 | X | X |
| Throw object | X | 9 | X, 1, 5 | 8 |
| Touch | X, 6 | X, 9 | X, 5, 6 | X, 8 |
| ***Bodily acts*** |  |  |  |  |
| Beg mouth-hand | 6 | X | 6 | X, |
| Beg mouth-mouth | X, 6 | X | X, 5, 6 | X, 8 |
| Bite | X, 6 | X, 9 | X, 2, 5, 6 | X, 8 |
| Bite attempt | X, 6 | X, 7 | X, 5, 6 | X, |
| Dangle | X, 6 | X, 9 | X, 5, 6 | X |
| Head-butt |  |  | X, 3, 5 | X |
| Head-stand | 6 |  | X, 1, 5, 6 | X |
| Kiss | X, 6 | X, 7 | X, 5, 6 | X, 8 |
| Look | X | X | X, 1 | X, 8 |
| Look back | 1, 6 |  | X, 1, 4, 6 | X |
| Peer | X, 6 | X | X, 5, 6 | X, 8 |
| Present body part | X, 6 | X | X, 5, 6 | X, 8 |
| Present object | X, 6 | X, 9 | X, 5, 6 | X |
| Rise up |  | 7 | X | 8 |
| Roll on back | X, 1, 6 |  | X, 6 |  |
| Rub body |  |  | X, 5 | X |
| Somersault | X, 6 |  | X, 6 |  |
| Spin | X |  |  |  |
| Spit |  |  | X, (1) |  |
| Throw self | X |  | X | X |
| ***Facial acts*** |  |  |  |  |
| Flapped lip |  |  | X, 1, 5 |  |
| Play face | X, 1, 6 | X, 7 | X, 1, 5,6 | X, 8 |
| Pout face | 6 | X, 7 | X, 5, 6 | 8 |

**Table S4** Communicative repertoire for presumed goal “Play/Affiliate” separately for mother-offspring and same-aged interactions (N = number of signal types, n = number of cases).

|  | **Mother-offspring** | | | | **Peers** | | | |
| --- | --- | --- | --- | --- | --- | --- | --- | --- |
| Comm. act | Bornean | | Sumatran | | Bornean | | Sumatran | |
|  | captive | wild | captive | wild | captive | wild | captive | wild |
| bite | 8 | 233 | 9 | 1 | 5 | 30 | 7 | 1 |
| bite attempt | 6 | 118 | 7 | 11 | 2 | 15 | 9 | 3 |
| embrace | 1 | 5 | 10 | 6 |  |  | 2 | 1 |
| fling |  | 2 | 1 | 7 |  |  | 10 | 8 |
| grab/hold onto | 17 | 455 | 29 | 49 | 11 | 92 | 28 | 31 |
| hand on | 16 | 146 | 9 | 30 |  | 13 | 5 | 5 |
| head-butt |  |  | 4 | 1 |  |  | 4 |  |
| head-stand |  |  |  |  |  |  |  | 1 |
| hit/kick | 3 |  | 7 | 1 | 1 |  | 20 |  |
| kiss | 1 | 56 | 12 | 2 |  |  | 3 |  |
| look at | 5 | 24 | 2 | 21 |  | 2 | 14 | 23 |
| look back at |  |  |  |  |  |  | 6 |  |
| loud scratch |  |  |  | 7 |  |  |  |  |
| peer |  |  |  | 3 |  |  | 6 | 2 |
| play face | 4 | 7 | 7 | 2 |  | 1 | 11 | 10 |
| poke |  | 30 | 6 |  |  |  | 9 | 3 |
| pout face |  | 10 | 2 |  |  |  | 2 |  |
| present body part |  | 33 |  | 3 |  | 21 |  | 4 |
| present object | 1 | 1 | 1 | 2 |  |  | 2 |  |
| pull | 12 | 71 | 9 | 9 | 3 | 5 | 46 | 16 |
| push | 5 | 5 | 1 | 8 |  |  | 5 | 4 |
| raise limb | 2 |  | 1 | 1 | 3 |  | 3 | 2 |
| reach | 5 | 4 | 12 | 13 | 4 | 1 | 25 | 24 |
| rub on rec |  |  |  |  |  |  | 7 | 8 |
| rise up | 1 |  |  |  |  |  |  |  |
| roll on back | 1 |  | 1 |  |  |  |  |  |
| rub body |  |  | 1 | 7 |  |  |  |  |
| somersault | 1 |  | 1 |  | 1 |  | 3 |  |
| spin | 1 |  |  |  | 1 |  |  |  |
| swing/dangle | 9 | 24 | 5 | 24 | 4 | 10 | 17 | 15 |
| throw object |  |  |  |  |  |  | 1 |  |
| throw self | 8 |  | 13 | 1 | 2 |  | 14 |  |
| tickle | 1 |  | 7 | 4 |  |  |  |  |
| touch | 6 | 235 | 18 | 39 | 8 | 270 | 16 | 37 |
| **n** | **114** | **1459** | **175** | **252** | **45** | **460** | **275** | **198** |
| **N** | **22** | **18** | **25** | **24** | **12** | **11** | **26** | **19** |

**Table S5** Effects of orang-utan species, research setting and control variables on number of observed communicative acts per individual (N = 70) derived using a LMM with a Gaussian error structure and identity link function. Significant effects (*P* < 0.05) are depicted in italics. This dataset included all individuals irrespective of sampling effort.

| **Repertoire size** | Estimate | SE | Chi Square | *P* |
| --- | --- | --- | --- | --- |
| Intercept | -0.578 | 0.151 | - | - |
| Species [Sumatran] | 0.07 | 0.066 | 1.325 | 0.250 |
| *Setting [wild]* | *-0.353* | *0.067* | *13.780* | *<0.001* |
| Age class [young imm.] | 0.17 | 0.09 | 3.692 | 0.055 |
| Age class [old imm.] | 0.188 | 0.114 | 2.825 | 0.093 |
| Sex [male] | -0.059 | 0.074 | 0.603 | 0.437 |
| *No. presumed goals* | *0.114* | *0.036* | *10.573* | *0.001* |
| *No. instances* | *0.557* | *0.047* | *82.771* | *<0.001* |

***Presumed goals of communicative acts across settings***

To enable the analyses of interaction outcomes and functional specificity, we first identified the major contexts of interactions. Across all settings, Play/affiliate was the presumed goal associated with the most types of communicative acts, with a mean (± SD) of 25.3 (± 5.6) and total N = 35, followed by Share food/object (mean ± SD = 16 ± 2.7, N = 20), Co-locomote (9.5 ± 1, N = 14), Stop action (9.3 ± 5.9; N = 15; not coded in captive Borneans), Groom (6.5 ± 3, N = 13), Move away (4.3 ± 2.3, N = 8, not assessed in captive Borneans), and Sexual contact (4.3 ± 2.3, N = 7, not assessed in captive Borneans). For full details of presumed goals in relation to communicative acts, orang-utan species and setting, see Tab 3.

**Table S6** Communicative acts associated with different presumed goals in relation to setting and species

|  |  | **Bornean** | | **Sumatran** | |  |  |
| --- | --- | --- | --- | --- | --- | --- | --- |
| **Presumed goal** | **All** | **captive** | **wild** | **captive** | **wild** | **mean** | **SD** |
| Play/affiliate | 35 | 24 | 18 | 31 | 28 | 25.3 | 5.6 |
| Share food/object | 20 | 14 | 15 | 15 | 20 | 16.0 | 2.7 |
| Stop action | 15 | N/A | 5 | 15 | 7 | 9.0 | 5.3 |
| Co-locomote | 14 | 9 | 9 | 9 | 11 | 9.5 | 1.0 |
| Groom | 13 | 10 | 4 | 8 | 4 | 6.5 | 3.0 |
| Move away | 8 | N/A | 3 | 7 | 3 | 4.3 | 2.3 |
| Sexual contact | 7 | N/A | 3 | 7 | 3 | 4.3 | 2.3 |

***Functional specificity of communicative acts across settings***

On average, functional specificity (i.e. proportion of use towards single interaction outcome) was 0.81 ± 0.19 across all setting. It somewhat lower in captive than wild individuals among Borneans (captive: 0.82 ± 0.2; wild: 0.88 ± 0.18), and higher among Sumatrans (captive: 0.82 ± 0.16; wild: 0.72 ± 0.22). In Sumatrans, average functional specificity of captivity-only acts (*N* = 8) was 0.88 ± 0.16, as opposed to 0.81 ± 0.16 for communicative acts used in both wild and captive settings (*N* = 29). In Borneans, average functional specificity of captivity-only (*N* = 3) and other acts (*N* = 27) was more similar (0.89 ± 0.19 and 0.87 ± 0.15, respectively). These descriptive results (the small sample prevented inferential analyses) suggest that Sumatran use more communicative acts of high functional specificity (i.e. those with tight meanings, Cartmill and Byrne [2010]) in captive than wild settings, which seems to be driven by those acts used only captivity (and thus, “invented”). Adopting the classification of Cartmill & Byrne (2010), for Bornean orang-utans we found that only 13 out of 23 (56.5%) types of communicative acts used in captive settings had tight meanings, versus 20 of the 24 (83.3%) used in the wild. In contrast, 26 of the 37 (70.3%) different communicative acts used by Sumatran orang-utans in captive settings had tight meanings, whereas only 17 of 30 (56.7%) in the wild did (see Tab. S8 for overview of tight, loose and ambiguous meanings in relation to species and setting).

**Tab. S7** Functional specificity of communicative acts in relation to orang-utan species and research setting, adopting the classification of Cartmill and Byrne (6).

|  | Bornean | | Sumatran | |
| --- | --- | --- | --- | --- |
|  | captive | wild | captive | wild |
| ambiguous | 1 | 1 | 0 | 6 |
| loose | 9 | 3 | 11 | 7 |
| tight | 13 | 20 | 26 | 17 |
| **Total** | **23** | **24** | **37** | **30** |

**Table S8** Information in study subjects and sample size (Bor = Bornean orang-utan, Sum = Sumatran orang-utan, Ad = adult, Im = Older immature, Dp = Younger immature).

| **No. subject** | **ID** | **Setting** | **Species** | **Group** | **Age class** | **Sex** | **Repertoire size** | **No. contexts** | **No. samples** |
| --- | --- | --- | --- | --- | --- | --- | --- | --- | --- |
| 1 | BAJ | captive | Bor | Apenheul | Dp | M | 9 | 4 | 66 |
| 2 | BUD | captive | Bor | Cologne | Im | M | 4 | 1 | 25 |
| 3 | CAJ | captive | Bor | Cologne | Ad | F | 7 | 4 | 43 |
| 4 | CIR | captive | Bor | Cologne | Dp | F | 14 | 4 | 91 |
| 5 | CIT | captive | Bor | Cologne | Dp | F | 12 | 4 | 61 |
| 6 | COR | captive | Bor | Cologne | Ad | F | 9 | 4 | 31 |
| 7 | KAW | captive | Bor | Apenheul | Im | M | 12 | 4 | 62 |
| 8 | MAN | captive | Bor | Munster | Ad | F | 6 | 4 | 37 |
| 9 | MIY | captive | Bor | Munster | Dp | M | 14 | 4 | 86 |
| 10 | NIA | captive | Bor | Munster | Im | F | 2 | 1 | 14 |
| 11 | SAR | captive | Bor | Munster | Ad | F | 5 | 1 | 20 |
| 12 | WAT | captive | Bor | Apenheul | Ad | F | 10 | 4 | 67 |
| 13 | BRU | captive | Sum | Munich | Ad | M | 2 | 2 | 11 |
| 14 | CAH | captive | Sum | Zurich | Ad | F | 15 | 6 | 82 |
| 15 | DJA | captive | Sum | Zurich | Ad | M | 6 | 3 | 22 |
| 16 | HAD | captive | Sum | Zurich | Im | M | 21 | 4 | 103 |
| 17 | ISO | captive | Sum | Munich | Im | F | 16 | 6 | 132 |
| 18 | JAH | captive | Sum | Munich | Ad | F | 13 | 5 | 58 |
| 19 | JOL | captive | Sum | Munich | Im | F | 13 | 5 | 70 |
| 20 | MAL | captive | Sum | Zurich | Im | M | 27 | 5 | 342 |
| 21 | MAT | captive | Sum | Munich | Ad | F | 11 | 5 | 70 |
| 22 | MIM | captive | Sum | Zurich | Dp | F | 22 | 4 | 208 |
| 23 | PAN | captive | Sum | Zurich | Dp | F | 21 | 4 | 199 |
| 24 | QUE | captive | Sum | Munich | Dp | M | 25 | 5 | 287 |
| 25 | QUI | captive | Sum | Munich | Dp | M | 18 | 4 | 101 |
| 26 | RIA | captive | Sum | Zurich | Dp | F | 14 | 3 | 99 |
| 27 | RON | captive | Sum | Munich | Dp | F | 21 | 3 | 197 |
| 28 | SIT | captive | Sum | Munich | Ad | F | 17 | 5 | 129 |
| 29 | TIM | captive | Sum | Zurich | Ad | F | 7 | 6 | 50 |
| 30 | XIR | captive | Sum | Zurich | Ad | F | 10 | 5 | 52 |
| 31 | JAN | wild | Bor | Tuanan | Dp | F | 19 | 4 | 320 |
| 32 | JUN | wild | Bor | Tuanan | Ad | F | 10 | 6 | 96 |
| 33 | CAK | wild | Bor | Tuanan | Dp | M | 13 | 4 | 124 |
| 34 | CIA | wild | Bor | Tuanan | Ad | F | 7 | 3 | 45 |
| 35 | DAN | wild | Bor | Tuanan | Im | M | 4 | 2 | 17 |
| 36 | DAR | wild | Bor | Tuanan | Dp | M | 17 | 4 | 330 |
| 37 | DES | wild | Bor | Tuanan | Ad | F | 8 | 4 | 144 |
| 38 | KEC | wild | Bor | Tuanan | Dp | M | 8 | 4 | 54 |
| 39 | KER | wild | Bor | Tuanan | Ad | F | 11 | 4 | 223 |
| 40 | KET | wild | Bor | Tuanan | Dp | M | 19 | 4 | 467 |
| 41 | KON | wild | Bor | Tuanan | Ad | F | 10 | 3 | 69 |
| 42 | MAW | wild | Bor | Tuanan | Im | F | 6 | 3 | 46 |
| 43 | MER | wild | Bor | Tuanan | Dp | M | 14 | 4 | 418 |
| 44 | MIL | wild | Bor | Tuanan | Ad | F | 11 | 6 | 119 |
| 45 | MIN | wild | Bor | Tuanan | Ad | F | 9 | 3 | 100 |
| 46 | MOB | wild | Bor | Tuanan | Dp | F | 18 | 3 | 390 |
| 47 | NAP | wild | Bor | Tuanan | Ad | M | 5 | 3 | 69 |
| 48 | TIN | wild | Bor | Tuanan | Ad | F | 7 | 5 | 59 |
| 49 | TUK | wild | Bor | Tuanan | Dp | M | 13 | 4 | 125 |
| 50 | ZAK | wild | Bor | Tuanan | Dp | F | 3 | 3 | 16 |
| 51 | FRI | wild | Sum | Suaq | Ad | F | 9 | 5 | 44 |
| 52 | AMO | wild | Sum | Suaq | Dp | M | 1 | 1 | 5 |
| 53 | BEO | wild | Sum | Suaq | Ad | M | 1 | 2 | 6 |
| 54 | CIN | wild | Sum | Suaq | Dp | F | 19 | 6 | 225 |
| 55 | CIS | wild | Sum | Suaq | Ad | F | 7 | 3 | 41 |
| 56 | EDE | wild | Sum | Suaq | Dp | F | 24 | 5 | 391 |
| 57 | ELL | wild | Sum | Suaq | Ad | F | 10 | 4 | 113 |
| 58 | FRA | wild | Sum | Suaq | Dp | M | 24 | 6 | 371 |
| 59 | KRO | wild | Sum | Suaq | Im | M | 7 | 2 | 45 |
| 60 | LIL | wild | Sum | Suaq | Ad | F | 8 | 4 | 43 |
| 61 | LIS | wild | Sum | Suaq | Ad | F | 6 | 4 | 34 |
| 62 | LOI | wild | Sum | Suaq | Dp | M | 20 | 5 | 164 |
| 63 | LUT | wild | Sum | Suaq | Dp | M | 20 | 5 | 313 |
| 64 | NIB | wild | Sum | Suaq | Ad | M | 1 | 1 | 6 |
| 65 | PEP | wild | Sum | Suaq | Dp | M | 9 | 3 | 38 |
| 66 | PIN | wild | Sum | Suaq | Ad | F | 2 | 2 | 13 |
| 67 | TIA | wild | Sum | Suaq | Ad | F | 3 | 3 | 22 |
| 68 | TOR | wild | Sum | Suaq | Dp | M | 10 | 3 | 92 |
| 69 | UNF | wild | Sum | Suaq | Ad | M | 1 | 1 | 6 |
| 70 | YUL | wild | Sum | Suaq | Im | F | 2 | 1 | 8 |

**Table S9** Information on variables coded using BORIS (S = signaller, R = recipient).

| **Variable and levels** | **Coding type** | **Description** |
| --- | --- | --- |
| **Presumed goal** | single-select | Apparent aim of the signaller and function of communicative act |
| food/object share |  | Share/hand over food item or object with signaller (S) |
| groom |  | Groom S |
| move away |  | Move away from S |
| play/affiliate |  | Play or physically affiliate |
| sexual contact |  | Engage in sexual contact |
| stop action |  | Stop a certain behaviour (e.g. begging) from R |
| joint travel |  | Initiate or coordinate joint travel |
| **Modality** | multi-select | Sensory modality in which the signal is perceived by the recipient |
| auditory |  | S's behaviour produces a sound and is perceived through hearing |
| visual |  | S's behaviour is perceived via eye sight |
| seismic |  | S's behaviour is perceived through substrate movement *(like vibration = indirect mechanic modality)* |
| tactile |  | S's behaviour is perceived through body contact |
| **Objects** | single-select | Objects involved by signaller in behaviour/interaction |
| none |  | No object involved |
| immobile |  | Immobile object involved, e.g. Tree branch that is still attached to tree |
| mobile |  | Mobile object involved, e.g. Loose branch, stone, stick, leaf…. |
| NA |  | Not clearly visible, unknown |
| **Response** | single-select | Response of recipient in reaction to S's behaviour |
| no reaction |  | No response or ignorance by R |
| ASO |  | "Apparently Satisfactory Outcome": R responds in a way that seems appropriate to S's behaviour |
| responds w/ signal |  | R responds with a signal - "negotiation" |
| visual attention |  | R looks at S (but no other response) |
| move away |  | R moves away from S |
| agonistic |  | R reacts in aggressive or agonistic way to S's behaviour |
| NA |  | Not clearly visible, unknown |

**Supplementary Figures**

Cumulative number of communicative atcs

| Number of observations | Apenheul | Cologne | Munster |
| --- | --- | --- | --- |
| 1 | 3 | 2 | 5 |
| 2 | 5 | 8 | 9 |
| 3 | 6 | 10 | 10 |
| 4 | 9 | 10 | 11 |
| 5 | 9 | 11 | 12 |
| 6 | 12 | 12 | 12 |
| 7 | 12 | 12 | 13 |
| 8 | 13 | 13 | 13 |
| 9 | 14 | 13 | 16 |
| 10 | 15 | 13 | 16 |
| 11 | 19 | 16 | 17 |
| 12 | 19 | 16 | 17 |
| 13 | 19 | 16 | 19 |
| 14 | 19 | 17 | 19 |
| 15 | 20 | 18 | 19 |
| 16 | 20 | 18 | 19 |
| 17 | 20 | 18 | 19 |
| 18 | 20 | 18 | 19 |
| 19 | 21 | 20 | 19 |
| 20 | 21 | 20 | 20 |
| 21 | 21 | 20 | 20 |
| 22 | 21 | 20 | 20 |
| 23 | 21 | 20 | 20 |
| 24 | 21 | 20 | 20 |
| 25 | 21 | 20 | 20 |
| 26 | 21 | 20 | 20 |
| 27 | 21 | 20 | 20 |
| 28 | 21 | 20 |  |
| 29 | 21 | 20 |  |
| 30 | 23 | 21 |  |
| 31 | 23 | 21 |  |
| 32 | 23 | 21 |  |
| 33 | 23 | 21 |  |
| 34 | 23 | 21 |  |
| 35 | 24 | 21 |  |
| 36 | 24 | 21 |  |
| 37 | 24 | 21 |  |
| 38 |  | 22 |  |
| 39 |  | 22 |  |
| 40 |  | 22 |  |
| 41 |  | 22 |  |
| 42 |  | 22 |  |
| 43 |  | 23 |  |
| 44 |  | 23 |  |
| 45 |  | 23 |  |
| 46 |  | 23 |  |
| 47 |  | 23 |  |
| 48 |  | 23 |  |
| 49 |  | 24 |  |
| 50 |  | 24 |  |
| 51 |  | 24 |  |
| 52 |  | 25 |  |
| 53 |  | 25 |  |
| 54 |  | 25 |  |
| 55 |  | 25 |  |
| 56 |  | 25 |  |

Cumulative number of communicative acts

| Number of observations | Hellabrunn | Zurich |
| --- | --- | --- |
| 1 | 5 | 1 |
| 2 | 5 | 2 |
| 3 | 7 | 3 |
| 4 | 8 | 4 |
| 5 | 9 | 4 |
| 6 | 10 | 7 |
| 7 | 10 | 8 |
| 8 | 11 | 8 |
| 9 | 11 | 9 |
| 10 | 11 | 9 |
| 11 | 11 | 12 |
| 12 | 11 | 12 |
| 13 | 12 | 13 |
| 14 | 13 | 13 |
| 15 | 14 | 14 |
| 16 | 15 | 14 |
| 17 | 15 | 16 |
| 18 | 16 | 17 |
| 19 | 16 | 17 |
| 20 | 16 | 18 |
| 21 | 16 | 18 |
| 22 | 16 | 18 |
| 23 | 16 | 19 |
| 24 | 17 | 19 |
| 25 | 17 | 19 |
| 26 | 17 | 20 |
| 27 | 17 | 21 |
| 28 | 19 | 21 |
| 29 | 19 | 21 |
| 30 | 19 | 22 |
| 31 | 19 | 23 |
| 32 | 19 | 23 |
| 33 | 19 | 23 |
| 34 | 19 | 23 |
| 35 | 19 | 23 |
| 36 | 19 | 23 |
| 37 | 19 | 23 |
| 38 | 19 | 23 |
| 39 | 19 | 24 |
| 40 | 19 | 24 |
| 41 | 19 | 24 |
| 42 | 19 | 24 |
| 43 | 19 | 24 |
| 44 | 19 | 24 |
| 45 | 19 | 24 |
| 46 | 19 | 24 |
| 47 | 20 | 24 |
| 48 | 20 | 24 |
| 49 | 21 | 24 |
| 50 | 21 | 24 |
| 51 | 21 | 24 |
| 52 | 21 | 24 |
| 53 | 22 | 24 |
| 54 | 22 | 24 |
| 55 | 22 | 24 |
| 56 | 22 | 24 |
| 57 | 22 | 24 |
| 58 | 22 | 24 |
| 59 | 23 | 24 |
| 60 | 23 | 24 |
| 61 | 23 | 24 |
| 62 | 23 | 24 |
| 63 | 23 | 24 |
| 64 | 23 | 24 |
| 65 | 23 | 24 |
| 66 | 23 | 24 |
| 67 | 23 | 24 |
| 68 | 23 | 24 |
| 69 | 23 | 24 |
| 70 | 23 | 24 |
| 71 | 23 | 24 |
| 72 | 23 | 24 |
| 73 | 23 | 24 |
| 74 | 23 | 24 |
| 75 | 23 | 24 |
| 76 | 23 | 24 |
| 77 | 23 | 24 |
| 78 | 23 | 24 |
| 79 | 23 | 24 |
| 80 | 23 | 24 |
| 81 | 23 | 24 |
| 82 | 23 | 25 |
| 83 | 23 | 25 |
| 84 | 23 | 25 |
| 85 | 24 | 25 |
| 86 | 25 | 25 |
| 87 | 25 | 25 |
| 88 | 25 | 25 |
| 89 | 25 | 25 |
| 90 | 25 | 25 |
| 91 | 25 | 25 |
| 92 | 26 | 25 |
| 93 | 26 | 25 |
| 94 | 26 | 25 |
| 95 | 26 | 26 |
| 96 | 26 | 26 |
| 97 | 26 | 27 |
| 98 | 26 | 28 |
| 99 | 26 | 28 |
| 100 | 26 | 28 |
| 101 | 26 | 28 |
| 102 | 26 | 28 |
| 103 | 26 | 28 |
| 104 | 27 | 28 |
| 105 | 27 | 28 |
| 106 | 27 | 28 |
| 107 | 27 | 28 |
| 108 | 27 | 28 |
| 109 | 27 | 28 |
| 110 | 27 | 28 |
| 111 | 28 | 28 |
| 112 | 28 | 28 |
| 113 | 28 | 28 |
| 114 | 28 | 28 |
| 115 | 28 | 28 |
| 116 | 28 | 28 |
| 117 | 28 | 28 |
| 118 | 29 | 28 |
| 119 | 29 | 28 |
| 120 | 29 | 28 |
| 121 | 29 | 28 |
| 122 | 30 | 28 |
| 123 | 30 | 28 |
| 124 | 30 | 29 |
| 125 | 30 | 29 |
| 126 | 30 | 29 |
| 127 | 30 | 29 |
| 128 | 30 | 29 |
| 129 | 30 | 29 |
| 130 | 30 | 29 |
| 131 | 30 | 29 |
| 132 | 30 | 29 |
| 133 | 30 | 29 |
| 134 | 30 | 29 |
| 135 | 30 | 29 |
| 136 | 30 | 29 |
| 137 | 30 | 29 |
| 138 | 30 | 31 |
| 139 | 30 | 31 |
| 140 | 30 | 31 |
| 141 | 30 | 31 |
| 142 | 30 | 31 |
| 143 | 31 | 31 |
| 144 | 31 | 31 |
| 145 | 31 | 31 |
| 146 | 33 | 31 |
| 147 | 33 | 31 |
| 148 | 33 | 31 |
| 149 | 33 | 31 |
| 150 | 33 | 31 |
| 151 | 33 | 31 |
| 152 | 33 | 31 |
| 153 | 33 | 32 |
| 154 | 33 | 32 |
| 155 | 33 | 32 |
| 156 | 33 | 32 |
| 157 | 33 | 32 |
| 158 | 33 | 32 |
| 159 | 33 | 32 |
| 160 | 33 | 32 |
| 161 | 33 | 32 |
| 162 | 33 | 32 |
| 163 | 33 | 32 |
| 164 | 33 | 32 |
| 165 | 33 | 32 |
| 166 | 33 | 32 |
| 167 | 34 | 32 |
| 168 | 34 | 32 |
| 169 | 34 | 32 |
| 170 | 34 | 32 |
| 171 | 34 | 32 |
| 172 | 34 | 32 |
| 173 | 34 | 33 |
| 174 | 34 | 33 |
| 175 | 34 | 33 |
| 176 | 34 | 33 |
| 177 | 34 | 33 |
| 178 | 34 | 33 |
| 179 | 34 | 33 |
| 180 | 34 | 33 |
| 181 | 34 | 33 |
| 182 | 34 | 33 |
| 183 | 34 | 33 |
| 184 | 34 | 33 |
| 185 | 35 | 33 |
| 186 | 35 | 33 |
| 187 | 35 | 33 |
| 188 | 35 | 33 |
| 189 | 35 | 33 |
| 190 | 35 | 33 |
| 191 | 35 | 33 |
| 192 | 35 | 33 |
| 193 | 35 | 33 |
| 194 | 35 | 33 |
| 195 | 35 | 33 |
| 196 | 35 | 33 |
| 197 | 35 | 33 |
| 198 | 35 | 33 |
| 199 | 35 | 33 |
| 200 | 35 | 33 |

**Figure S1** Cumulative number of identified communicative acts over observation time, depicted separately for captive study groups of Sumatran (left) and Bornean orang-utans (right).

Cumulative number of communicative acts

| Number of observations | Suaq | Tuanan |
| --- | --- | --- |
| 1 | 0 | 0 |
| 2 | 2 | 1 |
| 3 | 4 | 2 |
| 4 | 7 | 3 |
| 5 | 7 | 3 |
| 6 | 7 | 3 |
| 7 | 9 | 5 |
| 8 | 9 | 7 |
| 9 | 10 | 7 |
| 10 | 12 | 7 |
| 11 | 12 | 11 |
| 12 | 12 | 11 |
| 13 | 12 | 11 |
| 14 | 12 | 11 |
| 15 | 14 | 11 |
| 16 | 14 | 11 |
| 17 | 15 | 11 |
| 18 | 15 | 11 |
| 19 | 15 | 11 |
| 20 | 16 | 11 |
| 21 | 16 | 11 |
| 22 | 16 | 11 |
| 23 | 16 | 11 |
| 24 | 16 | 12 |
| 25 | 16 | 12 |
| 26 | 16 | 12 |
| 27 | 16 | 12 |
| 28 | 17 | 12 |
| 29 | 17 | 12 |
| 30 | 17 | 12 |
| 31 | 18 | 13 |
| 32 | 18 | 13 |
| 33 | 18 | 13 |
| 34 | 19 | 13 |
| 35 | 20 | 13 |
| 36 | 20 | 13 |
| 37 | 21 | 13 |
| 38 | 23 | 14 |
| 39 | 23 | 15 |
| 40 | 23 | 15 |
| 41 | 24 | 15 |
| 42 | 25 | 15 |
| 43 | 25 | 16 |
| 44 | 25 | 16 |
| 45 | 25 | 16 |
| 46 | 25 | 16 |
| 47 | 25 | 16 |
| 48 | 25 | 16 |
| 49 | 25 | 16 |
| 50 | 25 | 16 |
| 51 | 25 | 16 |
| 52 | 25 | 16 |
| 53 | 25 | 16 |
| 54 | 25 | 16 |
| 55 | 25 | 17 |
| 56 | 25 | 17 |
| 57 | 25 | 18 |
| 58 | 25 | 18 |
| 59 | 25 | 18 |
| 60 | 25 | 18 |
| 61 | 25 | 18 |
| 62 | 25 | 18 |
| 63 | 25 | 18 |
| 64 | 25 | 18 |
| 65 | 25 | 18 |
| 66 | 25 | 18 |
| 67 | 25 | 18 |
| 68 | 25 | 18 |
| 69 | 25 | 18 |
| 70 | 25 | 18 |
| 71 | 25 | 18 |
| 72 | 25 | 18 |
| 73 | 25 | 18 |
| 74 | 25 | 18 |
| 75 | 25 | 18 |
| 76 | 25 | 18 |
| 77 | 25 | 18 |
| 78 | 25 | 18 |
| 79 | 26 | 18 |
| 80 | 26 | 18 |
| 81 | 26 | 18 |
| 82 | 26 | 18 |
| 83 | 26 | 18 |
| 84 | 26 | 18 |
| 85 | 26 | 18 |
| 86 | 27 | 18 |
| 87 | 27 | 18 |
| 88 | 27 | 18 |
| 89 | 27 | 18 |
| 90 | 27 | 19 |
| 91 | 27 | 19 |
| 92 | 27 | 19 |
| 93 | 27 | 19 |
| 94 | 27 | 19 |
| 95 | 27 | 19 |
| 96 | 27 | 19 |
| 97 | 27 | 19 |
| 98 | 27 | 19 |
| 99 | 27 | 19 |
| 100 | 27 | 19 |
| 101 | 27 | 19 |
| 102 | 27 | 19 |
| 103 | 27 | 19 |
| 104 | 27 | 19 |
| 105 | 27 | 19 |
| 106 | 27 | 19 |
| 107 | 27 | 19 |
| 108 | 27 | 19 |
| 109 | 27 | 19 |
| 110 | 27 | 19 |
| 111 | 27 | 19 |
| 112 | 27 | 19 |
| 113 | 27 | 19 |
| 114 | 27 | 19 |
| 115 | 27 | 19 |
| 116 | 27 | 19 |
| 117 | 27 | 19 |
| 118 | 27 | 19 |
| 119 | 27 | 19 |
| 120 | 27 | 19 |
| 121 | 27 | 19 |
| 122 | 27 | 19 |
| 123 | 27 | 20 |
| 124 | 27 | 20 |
| 125 | 27 | 20 |
| 126 | 27 | 20 |
| 127 | 27 | 20 |
| 128 | 27 | 20 |
| 129 | 28 | 20 |
| 130 | 28 | 20 |
| 131 | 28 | 20 |
| 132 | 28 | 20 |
| 133 | 28 | 20 |
| 134 | 28 | 20 |
| 135 | 28 | 20 |
| 136 | 28 | 20 |
| 137 | 28 | 20 |
| 138 | 28 | 20 |
| 139 | 28 | 20 |
| 140 | 28 | 20 |
| 141 | 28 | 20 |
| 142 | 28 | 20 |
| 143 | 28 | 20 |
| 144 | 28 | 20 |
| 145 | 28 | 20 |
| 146 | 28 | 20 |
| 147 | 28 | 20 |
| 148 | 28 | 20 |
| 149 | 28 | 20 |
| 150 | 28 | 20 |
| 151 | 28 | 20 |
| 152 | 28 | 20 |
| 153 | 28 | 20 |
| 154 | 28 | 20 |
| 155 | 29 | 20 |
| 156 | 29 | 20 |
| 157 | 29 | 20 |
| 158 | 29 | 20 |
| 159 | 29 | 20 |
| 160 | 29 | 20 |
| 161 | 29 | 20 |
| 162 | 29 | 20 |
| 163 | 29 | 20 |
| 164 | 29 | 20 |
| 165 | 29 | 20 |
| 166 | 29 | 20 |
| 167 | 29 | 20 |
| 168 | 29 | 20 |
| 169 | 29 | 20 |
| 170 | 29 | 20 |
| 171 | 29 | 20 |
| 172 | 29 | 20 |
| 173 | 29 | 20 |
| 174 | 29 | 20 |
| 175 | 29 | 20 |
| 176 | 29 | 20 |
| 177 | 29 | 20 |
| 178 | 29 | 20 |
| 179 | 29 | 20 |
| 180 | 29 | 20 |
| 181 | 29 | 20 |
| 182 | 29 | 20 |
| 183 | 29 | 20 |
| 184 | 29 | 20 |
| 185 | 29 | 20 |
| 186 | 29 | 20 |
| 187 | 29 | 20 |
| 188 | 29 | 20 |
| 189 | 29 | 20 |
| 190 | 29 | 20 |
| 191 | 29 | 20 |
| 192 | 29 | 20 |
| 193 | 29 | 20 |
| 194 | 29 | 20 |
| 195 | 29 | 20 |
| 196 | 29 | 20 |
| 197 | 29 | 20 |
| 198 | 29 | 20 |
| 199 | 29 | 20 |
| 200 | 29 | 20 |

**Figure S2** Cumulative number of identified communicative acts over observation time, depicted separately for wild populations of Sumatran (Suaq) and Bornean (Tuanan) orang-utans.

**Figure S3** Cumulative number of identified communicative acts over observation time, depicted separately for three highly sampled individuals from different study groups.

Ketambe (Tuanan)

| Number of consecutive observations | Cumulative number of comm. acts |
| --- | --- |
| 1 | 1 |
| 2 | 2 |
| 3 | 6 |
| 4 | 7 |
| 5 | 7 |
| 6 | 8 |
| 7 | 9 |
| 8 | 9 |
| 9 | 9 |
| 10 | 9 |
| 11 | 9 |
| 12 | 9 |
| 13 | 9 |
| 14 | 10 |
| 15 | 11 |
| 16 | 11 |
| 17 | 11 |
| 18 | 11 |
| 19 | 11 |
| 20 | 11 |
| 21 | 11 |
| 22 | 11 |
| 23 | 11 |
| 24 | 11 |
| 25 | 12 |
| 26 | 12 |
| 27 | 13 |
| 28 | 13 |
| 29 | 13 |
| 30 | 13 |
| 31 | 13 |
| 32 | 13 |
| 33 | 13 |
| 34 | 13 |
| 35 | 13 |
| 36 | 13 |
| 37 | 13 |
| 38 | 13 |
| 39 | 14 |
| 40 | 14 |
| 41 | 15 |
| 42 | 16 |
| 43 | 16 |
| 44 | 16 |
| 45 | 16 |
| 46 | 16 |
| 47 | 16 |
| 48 | 17 |
| 49 | 17 |
| 50 | 17 |
| 51 | 17 |
| 52 | 17 |
| 53 | 17 |
| 54 | 17 |
| 55 | 17 |
| 56 | 17 |
| 57 | 17 |
| 58 | 17 |
| 59 | 17 |
| 60 | 17 |
| 61 | 17 |
| 62 | 17 |
| 63 | 17 |
| 64 | 17 |
| 65 | 17 |
| 66 | 17 |
| 67 | 17 |
| 68 | 17 |
| 69 | 17 |
| 70 | 17 |
| 71 | 17 |
| 72 | 17 |
| 73 | 17 |
| 74 | 17 |
| 75 | 17 |
| 76 | 17 |
| 77 | 17 |
| 78 | 17 |
| 79 | 17 |
| 80 | 17 |
| 81 | 17 |
| 82 | 17 |
| 83 | 17 |
| 84 | 17 |
| 85 | 17 |
| 86 | 17 |
| 87 | 17 |
| 88 | 17 |
| 89 | 17 |
| 90 | 17 |
| 91 | 17 |
| 92 | 17 |
| 93 | 17 |
| 94 | 17 |
| 95 | 17 |
| 96 | 17 |
| 97 | 17 |
| 98 | 17 |
| 99 | 17 |
| 100 | 17 |
| 101 | 17 |
| 102 | 17 |
| 103 | 17 |
| 104 | 17 |
| 105 | 17 |
| 106 | 17 |
| 107 | 17 |
| 108 | 17 |
| 109 | 17 |
| 110 | 18 |
| 111 | 18 |
| 112 | 18 |
| 113 | 18 |
| 114 | 18 |
| 115 | 18 |
| 116 | 18 |
| 117 | 18 |
| 118 | 18 |
| 119 | 18 |
| 120 | 18 |
| 121 | 18 |
| 122 | 18 |
| 123 | 18 |
| 124 | 18 |
| 125 | 18 |
| 126 | 18 |
| 127 | 18 |
| 128 | 18 |
| 129 | 18 |
| 130 | 18 |
| 131 | 18 |
| 132 | 18 |
| 133 | 18 |
| 134 | 18 |
| 135 | 18 |
| 136 | 18 |
| 137 | 18 |
| 138 | 18 |
| 139 | 18 |
| 140 | 18 |
| 141 | 18 |
| 142 | 18 |
| 143 | 18 |
| 144 | 18 |
| 145 | 18 |
| 146 | 18 |
| 147 | 18 |
| 148 | 18 |
| 149 | 18 |
| 150 | 18 |

Malou (Zurich)

| Number of consecutive observations | Cumulative number of comm. acts |
| --- | --- |
| 1 | 1 |
| 2 | 1 |
| 3 | 3 |
| 4 | 4 |
| 5 | 4 |
| 6 | 6 |
| 7 | 6 |
| 8 | 7 |
| 9 | 9 |
| 10 | 10 |
| 11 | 11 |
| 12 | 13 |
| 13 | 14 |
| 14 | 14 |
| 15 | 14 |
| 16 | 15 |
| 17 | 16 |
| 18 | 17 |
| 19 | 17 |
| 20 | 17 |
| 21 | 18 |
| 22 | 18 |
| 23 | 18 |
| 24 | 18 |
| 25 | 18 |
| 26 | 18 |
| 27 | 18 |
| 28 | 18 |
| 29 | 19 |
| 30 | 19 |
| 31 | 19 |
| 32 | 19 |
| 33 | 19 |
| 34 | 19 |
| 35 | 20 |
| 36 | 20 |
| 37 | 21 |
| 38 | 21 |
| 39 | 21 |
| 40 | 21 |
| 41 | 22 |
| 42 | 22 |
| 43 | 22 |
| 44 | 22 |
| 45 | 22 |
| 46 | 22 |
| 47 | 22 |
| 48 | 22 |
| 49 | 22 |
| 50 | 23 |
| 51 | 24 |
| 52 | 24 |
| 53 | 26 |
| 54 | 26 |
| 55 | 26 |
| 56 | 26 |
| 57 | 26 |
| 58 | 26 |
| 59 | 26 |
| 60 | 26 |
| 61 | 26 |
| 62 | 26 |
| 63 | 26 |
| 64 | 26 |
| 65 | 26 |
| 66 | 26 |
| 67 | 26 |
| 68 | 26 |
| 69 | 26 |
| 70 | 26 |
| 71 | 26 |
| 72 | 26 |
| 73 | 26 |
| 74 | 26 |
| 75 | 26 |
| 76 | 26 |
| 77 | 26 |
| 78 | 26 |
| 79 | 26 |
| 80 | 26 |
| 81 | 26 |
| 82 | 27 |
| 83 | 27 |
| 84 | 27 |
| 85 | 28 |
| 86 | 28 |
| 87 | 28 |
| 88 | 28 |
| 89 | 28 |
| 90 | 29 |
| 91 | 29 |
| 92 | 29 |
| 93 | 29 |
| 94 | 29 |
| 95 | 29 |
| 96 | 29 |
| 97 | 29 |
| 98 | 29 |
| 99 | 29 |
| 100 | 29 |
| 101 | 29 |
| 102 | 29 |
| 103 | 29 |
| 104 | 29 |
| 105 | 29 |
| 106 | 29 |
| 107 | 29 |
| 108 | 29 |
| 109 | 29 |
| 110 | 29 |
| 111 | 29 |
| 112 | 29 |
| 113 | 29 |
| 114 | 29 |
| 115 | 29 |
| 116 | 29 |
| 117 | 29 |
| 118 | 29 |
| 119 | 29 |
| 120 | 29 |
| 121 | 29 |
| 122 | 29 |
| 123 | 29 |

Frankie (Suaq)

| Number of consecutive observations | Cumulative number of comm. acts |
| --- | --- |
| 1 | 3 |
| 2 | 6 |
| 3 | 6 |
| 4 | 6 |
| 5 | 6 |
| 6 | 10 |
| 7 | 10 |
| 8 | 10 |
| 9 | 11 |
| 10 | 11 |
| 11 | 11 |
| 12 | 11 |
| 13 | 11 |
| 14 | 11 |
| 15 | 11 |
| 16 | 12 |
| 17 | 13 |
| 18 | 13 |
| 19 | 14 |
| 20 | 14 |
| 21 | 14 |
| 22 | 15 |
| 23 | 16 |
| 24 | 16 |
| 25 | 18 |
| 26 | 19 |
| 27 | 19 |
| 28 | 20 |
| 29 | 21 |
| 30 | 21 |
| 31 | 21 |
| 32 | 21 |
| 33 | 21 |
| 34 | 21 |
| 35 | 21 |
| 36 | 22 |
| 37 | 23 |
| 38 | 23 |
| 39 | 24 |
| 40 | 24 |
| 41 | 24 |
| 42 | 24 |
| 43 | 24 |
| 44 | 24 |
| 45 | 24 |
| 46 | 24 |
| 47 | 24 |
| 48 | 24 |
| 49 | 24 |
| 50 | 24 |
| 51 | 24 |
| 52 | 24 |
| 53 | 24 |
| 54 | 24 |
| 55 | 24 |
| 56 | 24 |
| 57 | 24 |
| 58 | 24 |
| 59 | 24 |
| 60 | 24 |
| 61 | 24 |
| 62 | 24 |
| 63 | 24 |
| 64 | 24 |
| 65 | 24 |
| 66 | 24 |
| 67 | 24 |
| 68 | 24 |
| 69 | 24 |
| 70 | 24 |
| 71 | 24 |
| 72 | 24 |
| 73 | 24 |
| 74 | 24 |
| 75 | 24 |
| 76 | 24 |
| 77 | 24 |
| 78 | 24 |
| 79 | 24 |
| 80 | 24 |
| 81 | 25 |
| 82 | 25 |
| 83 | 25 |
| 84 | 25 |
| 85 | 25 |
| 86 | 25 |
| 87 | 25 |
| 88 | 25 |
| 89 | 26 |
| 90 | 26 |
| 91 | 26 |
| 92 | 26 |
| 93 | 26 |
| 94 | 26 |
| 95 | 26 |
| 96 | 26 |
| 97 | 26 |
| 98 | 27 |
| 99 | 27 |
| 100 | 27 |
| 101 | 27 |
| 102 | 27 |
| 103 | 27 |
| 104 | 27 |
| 105 | 27 |
| 106 | 27 |
| 107 | 27 |
| 108 | 27 |
| 109 | 27 |
| 110 | 27 |
| 111 | 27 |
| 112 | 27 |
| 113 | 27 |
| 114 | 27 |
| 115 | 27 |
| 116 | 27 |
| 117 | 27 |
| 118 | 27 |
| 119 | 27 |
| 120 | 27 |
| 121 | 27 |
| 122 | 27 |
| 123 | 27 |
| 124 | 27 |
| 125 | 27 |
| 126 | 27 |
| 127 | 27 |
| 128 | 27 |

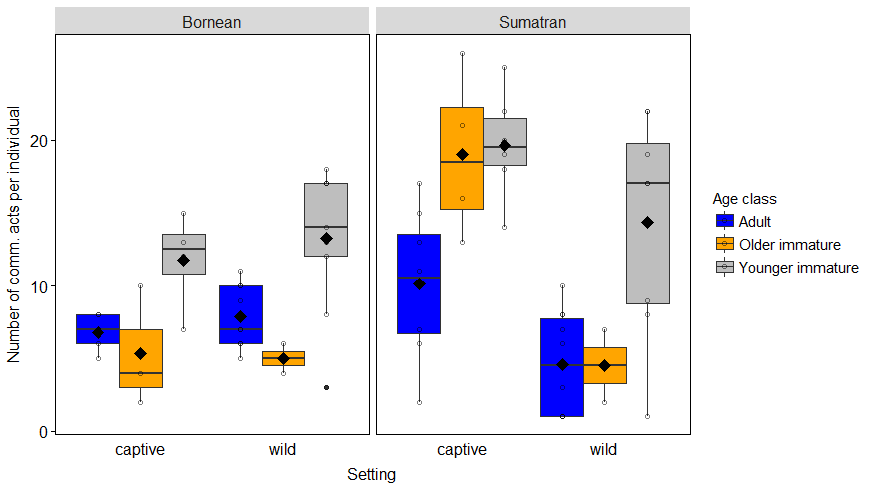
**Figure S4** Number of identified communicative acts in relation to orang-utan species, research setting and age class, using the entire dataset of 70 individuals. Indicated are individual means (circles), population means (filled diamonds), medians (horizontal lines), quartiles (boxes), percentiles (2.5% and 97.5%, vertical lines) and outliers (filled dots).
